# CHAMPOLLION: Robust Multi-Omics Integration via Inverse Optimal Transport Using Paired Cells

**DOI:** 10.64898/2026.04.28.721317

**Authors:** Jules Samaran, Gabriel Peyré, Laura Cantini

## Abstract

Fully capturing cellular identity requires integrating multiple molecular layers. Bridge integration, i.e. aligning unimodal datasets using a paired multi-omic reference, has emerged as a practical solution, yet existing methods offer limited interpretability and use paired information without regularization, making them sensitive to limited size and coverage. We introduce CHAMPOLLION, which uses regularized optimal transport to learn an interpretable cross-modal metric that drives the alignment of unpaired cells while capturing relationships between molecular features. Benchmarks on RNA-protein and RNA-ATAC datasets show that CHAMPOLLION outperforms existing approaches, remaining accurate with few paired cells and even generalizing to unseen cell types. Beyond alignment, CHAMPOLLION reveals biologically meaningful cross-modal relationships, highlighting in scRNA-protein data a potential role for CD18 across multiple cancers, and, in a human tonsil atlas combining scRNA-seq and scATAC-seq, suggesting that MEF2C may regulate inflammatory responses beyond the brain, notably in plasmacytoid dendritic cells.

## 1 Introduction

Over the past decade single-cell sequencing has expanded far beyond its scRNA-seq origins [1]. High-throughput protocols now capture chromatin accessibility [2], DNA methylation [3], protein abundance [4] and other layers at single-cell resolution, and recent “multi-omic” assays such as CITE-seq [5], TEA-seq [6] or the 10x Multiome platform can even profile two or three of these modalities in the same cells. Because each layer reveals a different facet of cellular regulation, jointly analyzing them offers a more complete view of cell identity than any modality alone. In principle, paired multi-omic datasets provide a perfect opportunity to uncover cellular identities by integrating multiple modalities, and a growing toolbox has been developed to analyze them [7]. In practice, however, such datasets rarely cover more than two modalities, and typically trade depth for breadth: per-cell read counts are lower, yielding sparsier measurements and lower library complexity than in dedicated single-omic experiments. For example, 10X Multiome yields about three-times fewer UMIs per 100 reads than a standalone 10x scRNA-seq run [8]. Large, high-quality unimodal collections therefore remain indispensable for resolving differences between closely related and rare cell populations.

To leverage this type of data, the community has therefore turned to diagonal integration, which seeks to align unpaired single-cell datasets that share neither cells nor measured features. Its precise aim is to embed them in a joint latent space where cells group by biology rather than by modality. The challenge is that each modality captures a distinct set of features; lacking a common reference, their profiles cannot be directly compared, making it easy to align cells incorrectly [9]. Sparse prior connections between features across modalities, such as mapping ATAC peaks to nearby genes or pairing transcripts with their encoded proteins, provide a common ground for comparing cells across assays. Consequently, every state-of-the-art diagonal integration method [10–12] incorporates this kind of prior knowledge to steer the alignment and avoid arbitrary overlays.

While these priors certainly aid alignment, profiling even a modest set of cells in both modalities (commonly called the *bridge* [13]) provides a direct anchor that can further orient and integrate larger unimodal datasets. This scenario is extremely relevant in practice: a lab might generate a small, application-specific multi-omic dataset and then complement it with deeper single-omic experiments, thereby retaining single-omic accuracy while still achieving cross-modal alignment through the paired dataset. Leveraging such a configuration is known as “bridge integration” and several recent methods exploit this idea [13–15]. Bridge datasets range from large, diverse references sufficient to align the data on their own to small, cell type limited subsets; consequently, an effective method must work across this spectrum and allow adjusting how heavily it relies on the paired cells. The best-performing bridge integration tools to date are Seurat’s dictionary-learning workflow [13] and a family of variational-autoencoder (VAE) models [15–18] whose scalable, modular designs can handle arbitrary patterns of missing modalities. However, all of these typically couple modalities solely through paired cells and ignore existing feature-level biological links, hampering performance when the bridge is small or does not cover all relevant cell populations. More generally, they lack regularizing mechanisms to control the influence of the bridge, making them sensitive to biases and missing data. Besides, they offer no interpretability of the inferred cross-modal relationships, making it difficult to extract biological insight beyond cell alignment. Some hybrid frameworks try to leverage prior links by imputing the missing modality [19–21], but they treat these predicted values exactly like true measurements, although the two differ appreciably. Lacking any mechanism to downplay the influence of imputed entries leaves the core challenge unresolved: the algorithm can over-trust noisy imputations and fail to capitalize on the rich bridge data.

Here we introduce CHAMPOLLION, a bridge-integration framework that uses inverse optimal transport [22–25] to learn a cross-modal metric from the paired measurements and then applies this metric to align unpaired cells. By jointly leveraging the paired data and informative feature-level priors, CHAMPOLLION is resilient to small or incomplete bridges, while an explicit regularization controls the relative influence of the bridge and mitigates biases arising from incomplete or unbalanced paired data. Finally, the metric learned by CHAMPOLLION is interpretable, and can thus offer mechanistic insights, revealing associations between features across modalities.

Here we introduce CHAMPOLLION, a bridge-integration framework that uses inverse optimal transport [22] to learn a cross-modal metric from paired measurements and apply it to align unpaired cells. By jointly leveraging paired data and feature-level priors, CHAMPOLLION remains robust to small or incomplete bridges, while an explicit regularization controls the relative influence of the bridge and mitigates biases arising from incomplete or unbalanced paired data. Importantly, the learned metric is interpretable, revealing how features from one modality correspond to and interact with those of another.

We benchmark CHAMPOLLION against existing state-of-the-art integration methods on scRNA-surface protein (Cite-seq) and scRNA-scATAC (10X Multiome) gold-standard datasets. Our benchmark includes different scenarios, varying the size of the bridge and its cell type coverage to challenge the methods. We show that CHAMPOLLION obtains the best performance in all settings, extracting maximal information from the bridge while its regularization provides robustness when that bridge is sparse or incomplete. Furthermore, we demonstrate the biological relevance of CHAMPOLLION through two case studies. In RNA-protein data, the learned cross-modal cost captures meaningful gene-protein relationships, notably recovering known pairs, but also enables discovery by suggesting novel cell type markers as well as functional insights for less-characterized proteins, including a potential role for CD18 in tumor progression across multiple cancers. In a RNA-ATAC human tonsil atlas, CHAMPOLLION shows that it can accurately integrate very large datasets while enabling mechanistic insights, notably suggesting that MEF2C acts as a broader regulator of inflammatory responses beyond the brain, including in plasmacytoid dendritic cells.

In practice, CHAMPOLLION is an efficient tool, leveraging both GPU acceleration and symbolic tensorial operations to scale to large datasets. It is implemented as a Python package seamlessly integrated within the scverse ecosystem and is available at https://github.com/cantinilab/champollion.

## 2 Results

### 2.1 CHAMPOLLION: a new tool for bridge integration

We developed CHAMPOLLION, a method for single-cell bridge integration.

As shown in Figure 1a, CHAMPOLLION considers two modalities for integration, labeled 1 and 2, and uses single-cell matrices as input. The paired measurements or “bridge” are 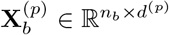 with *p* = 1, 2, measured on the same *n*_*b*_ cells, while the larger unimodal measurements which must be integrated are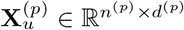, measured on disjoint sets of cells (therefore unpaired). We denote by 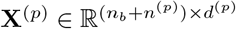 the full dataset obtained by concatenating bridge and unpaired cells along the cell dimension, 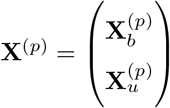. Leveraging feature-level priors, e.g. mapping ATAC peaks to their proximal genes, each input matrix **X**^(*p*)^ can be transformed into a shared feature space, yielding 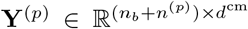, which provides a common basis for comparing cells across modalities. CHAMPOLLION is an Optimal Transport (OT) framework which leverages all these inputs to compute a cell-to-cell matching for the unpaired data which can then be used for visualization, transferring annotations across modalities and discovering subpopulations of cells.

**Fig. 1:**
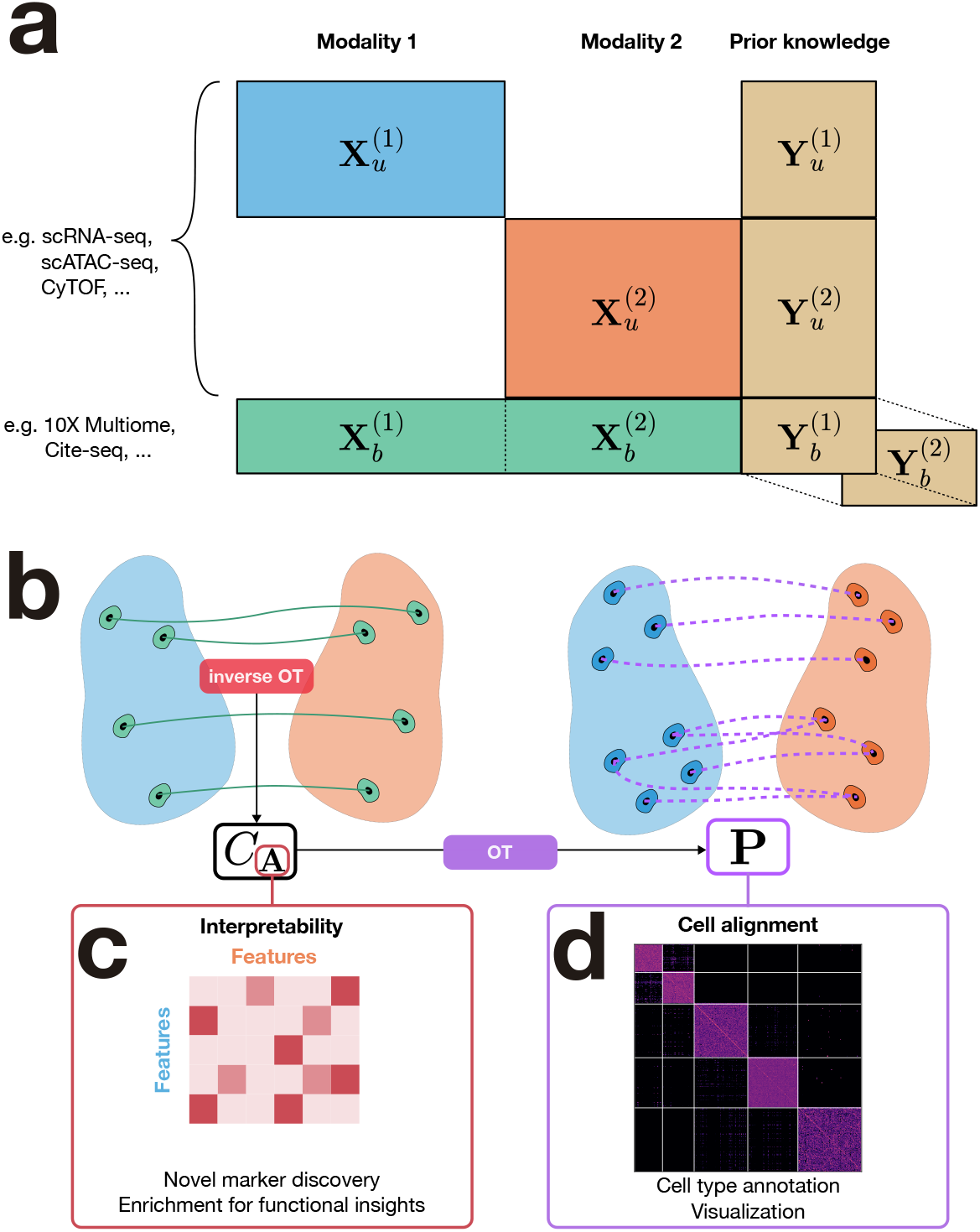
Overview of the CHAMPOLLION framework. **(a)** Schematic representation of the single-cell data used as input by CHAMPOLLION. For each modality (*p* = 1, 2), unpaired measurements 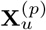 and a bridge 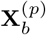 composed of paired cell measurements are available. Feature-level priors are leveraged to construct additional representations 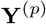 in a shared feature space, enabling direct cross-modal comparison; for bridge cells, 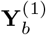 and 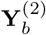 correspond to the same cells but are derived from each modality. **(b)** CHAMPOLLION proceeds in two steps: solve an inverse optimal transport problem on paired data to infer a cross-modal cost, which is then used to solve a standard optimal transport problem and align unpaired cells. **(c)** This yields an interpretable cross-modal cost parametrized by the matrix **A**, which captures feature-to-feature interactions across modalities and can be investigated for biological insights. **(d)** It also produces a transport plan between unpaired cells, enabling cross-modal information transfer, such as cell type annotation transfer, and joint visualization.

OT [26] has become a staple in machine learning and single-cell genomics for comparing probability distributions (here, populations of cells) by using a ground-cost matrix of pairwise cell distances to compute a transport plan, i.e. a probabilistic coupling between two datasets. Several cell-matching methods [27, 28] already use OT on the shared-feature space **Y**^(*p*)^, but classical OT cannot leverage paired samples. CHAMPOLLION overcomes this limitation with *inverse* OT [22–25]: starting from a known transport plan, here the paired bridge cells, it learns the underlying distance (ground-cost) function that would make that coupling optimal. As shown in Figure 1b, we thus first learn a parameterized ground cost *C*_**A**_ that best explains the paired data matrices 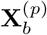 while also relying on the shared feature representations 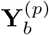. We then apply this learned cost to the 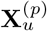 and 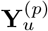 matrices and solve a standard OT problem, yielding a transport plan that matches unpaired cells across modalities.

In more detail, we define a parameterized cross-modal ground cost: 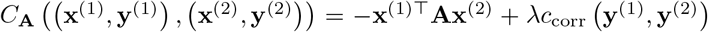 where **x**^(*p*)^ (respectively **y**^(*p*)^) is a single cell omics profile corresponding to a row of the complete **X**^(*p*)^ (respectively **Y**^(*p*)^) matrix, *c*_corr_ is a correlation-based dissimilarity, and the matrix **A** captures feature-to-feature interactions across modalities, with the negative sign ensuring that positive entries in **A** correspond to positive associations between features. We learn **A** by solving the inverse OT problem on the paired cells, adding an *ℓ*_1_ penalty ℛ (**A**) = *γ*∥**A**∥_1_ to encourage sparsity for interpretability (see Methods). CHAMPOLLION therefore delivers three outputs useful for downstream analyses (Figure 1c-d): (i) the interaction matrix **A**, which can highlight cross-modal regulatory links (e.g., protein influences on transcript levels); (ii) the learned cost function *C*_**A**_, enabling the computation of distances between any two cells from different modalities; and (iii) the OT transport plan that matches the previously unpaired cells.

The *ℓ*_1_ regularization on **A** serves two complementary purposes. First, by driving many interaction weights to zero it yields a sparse, more interpretable matrix in which the remaining entries stand out as putative cross-modal mechanistic links. Second, sparsity guards against overfitting when the bridge is small: with only a few paired cells, an unconstrained matrix could memorize noise. In effect, CHAMPOLLION blends two information sources: the weak feature-level prior carried by the **Y**^(*p*)^ measurements and the stronger, but potentially limited, paired cells in the bridge, allowing each to compensate for gaps in the other and to generalize to unseen cell types. The trade-off is tunable: raising *γ* shrinks **A**, reducing reliance on the paired samples and gradually reverting the cost to the prior alone, whereas lowering *γ* lets the model learn richer interactions when the bridge is large and reliable.

From a practical standpoint CHAMPOLLION is light-weight and flexible. Users may feed the **X**^(*p*)^ matrices with either raw single-cell profiles or any low-dimensional embeddings produced by other tools. CHAMPOLLION relies on GPU-accelerated algorithms that require no favorable initialization and run with a small memory footprint thanks to symbolic tensor operations [29]. The code is available as an scverse-compatible [30] Python package, so it plugs directly into existing preprocessing and visualization workflows.

We carried out extensive benchmarks against the current state-of-the-art. Although numerous methods have been proposed, we concentrate on Seurat [13] and MIDAS [16], which are both the most widely used and consistently top-performing tools for bridge integration [31].

### 2.2 CHAMPOLLION generalizes well to cell types absent from the paired dataset

Bridge integration relies on a small multi-omic “bridge” to align much larger single-omic datasets, so by design it must squeeze maximal information from only a few paired cells. The smaller that bridge, the greater the odds that certain cell populations, especially rare or condition-specific ones, will never be sampled. A viable bridge-integration strategy must therefore not only use the paired data efficiently but also generalize to cell types that are entirely absent from the bridge, a challenge that routinely arises in practice.

We assessed this aspect on a 10X Multiome human PBMC dataset (see Figure 2a), a well-annotated fully paired RNA+ATAC dataset from which we kept five populations: memory CD4 T cells, naive CD8 T cells, classical monocytes, and two closely related natural-killer subtypes: CD56^+^ and CD56^−^. Although every cell in the dataset comes with both modalities profiled and a cell type label, we purposely mask the RNA-ATAC pairing for a subset of cells to mimic an unpaired scenario and withhold all cell type annotations from the integration algorithms using the hidden pairings and labels solely for downstream evaluation. Half of the CD4 T, CD8 T and monocyte cells were assigned to the paired dataset, while the remaining cells (including *all* NK cells) were placed in the pseudo-unpaired set. In this design the integrator must extrapolate the alignment to cell populations absent from the bridge and, additionally, distinguish the two NK sub-populations, providing a stringent test of generalization.

**Fig. 2:**
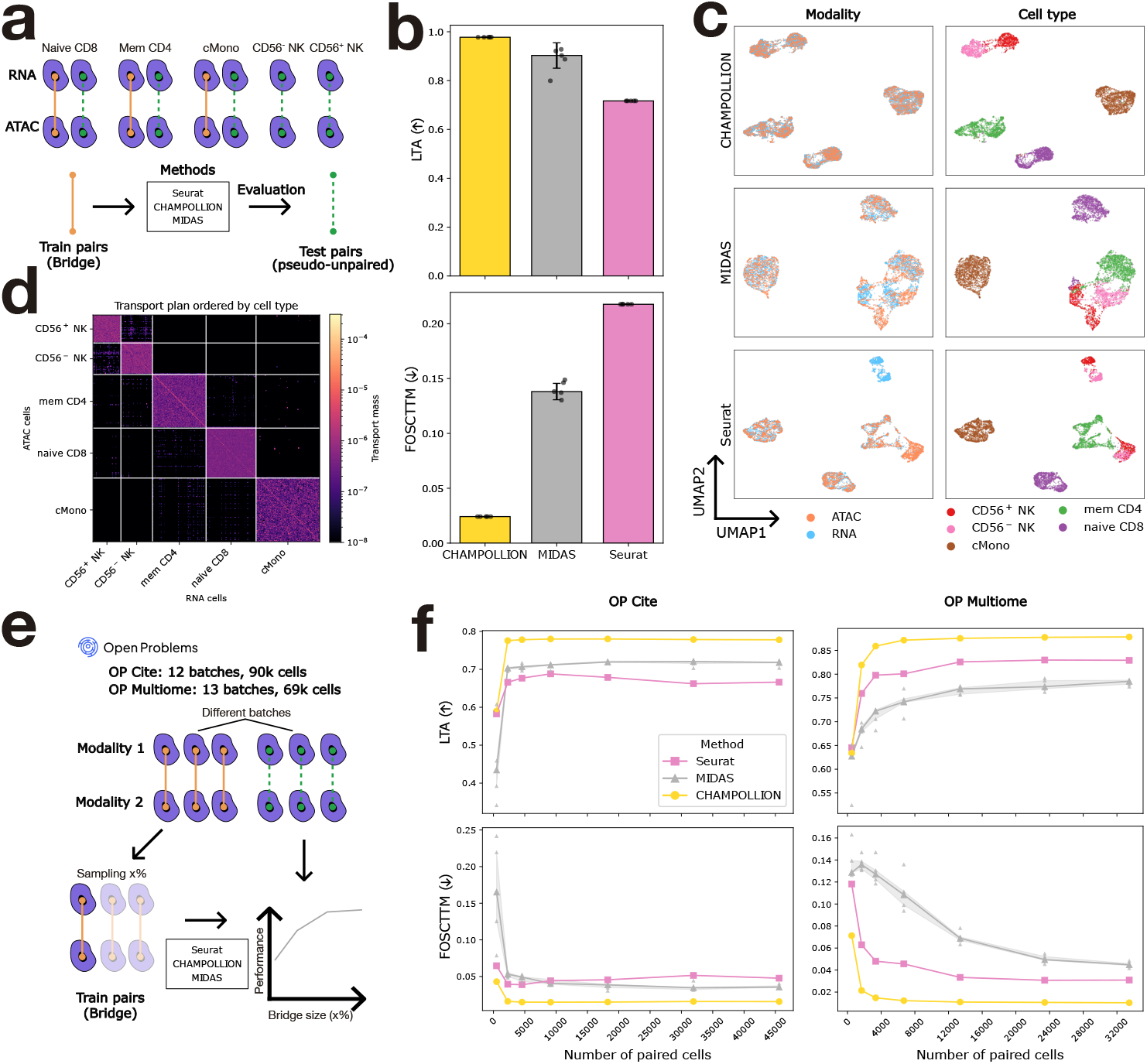
Benchmarking CHAMPOLLION against the state-of-the-art. **(a)** Schematic representation of the first benchmark experiment. A 10X Multiome dataset is split into a bridge and a test set of pseudo-unpaired cells, with the bridge containing only 3 of the 5 cell types present in the pseudo-unpaired set. **(b)** LTA (higher is better) and FOSCTTM (lower is better) scores for CHAMPOLLION, MIDAS, and Seurat on the setting described in (a). Bars show the median across *n* = 5 random initialization seeds, with error bars indicating standard deviation; dots correspond to individual runs. **(c)** UMAP visualizations of the pseudo-unpaired cells for each method. Cells are colored by modality (left) and by cell type (right). **(d)** Transport plan inferred by CHAMPOLLION for pseudo-unpaired cells. Rows and columns are ordered by cell type, and the diagonal corresponds to the true pairs between RNA and ATAC profiles. **(e)** Schematic representation of the second benchmark experiment. For two paired datasets (one Multiome and one CITE-seq), half of the cells are held out as the pseudo-unpaired test set, and increasing fractions of the remainder are used as the bridge to assess the impact of its size on performance. **(f)** LTA and FOSCTTM scores are reported on both datasets across bridge sizes as described in (e). Lines show median performance across *n* = 5 seeds, while shaded regions indicate the interquartile range, and points denote individual seed results.

To place our matching algorithm and the embedding-based baselines on comparable footing, we convert every output into a cross-modal similarity matrix: for the baselines we compute pairwise distances between their RNA and ATAC embeddings, whereas for our method we simply use the optimal transport plan itself. We then quantify biological fidelity with two standard, complementary metrics: (i) Label-Transfer Accuracy (LTA) [32], which measures how accurately cell type annotations are transferred from one modality to the other through the inferred similarities (range 0-1, higher is better), and (ii) Fraction Of Samples Closer Than the True Match (FOSCTTM) [33], which, for each cell, reports the proportion of opposite-modality cells that are nearer than its true pair, thus directly gauging pairwise alignment quality (range 0-1, lower is better).

Across both evaluation metrics CHAMPOLLION consistently outperformed all other methods, highlighting its robustness and capacity to generalize (see Figure 2b). CHAMPOLLION achieved an LTA of 0.977, indicating near-perfect alignment at the cell-type level, and an FOSCTTM of 0.024, reflecting accurate matching at single-cell resolution. Because MIDAS depends on randomly initialized neural networks with a non-convex objective, its scores varied across runs, unlike Seurat and CHAMPOLLION. While our approach aligns cells without embedding them in a common latent space, it is possible to use our inferred transport plan to project cells of one modality onto the other modality (see Methods). We were thus able to obtain UMAP visualizations for all methods which qualitatively mirrored the previous quantitative results (see Figure 2c). Whereas the baselines integrate the bridge-supported cell types nearly perfectly, they struggle to align the two NK subsets, with Seurat performing worst. In contrast, CHAMPOLLION not only aligns NK cells but also correctly distinguishes the CD56^+^ and CD56^−^ subpopulations. Beyond UMAP visualizations, the inferred transport plan exhibited a clear block-diagonal structure, reflecting the high LTA achieved by CHAMPOLLION (see Figure 2d). Within each block, a finer diagonal pattern indicates preferential matching to true counterparts rather than to cells of the same type, consistent with the low FOSCTTM.

### 2.3 Superior performance in scRNA-surface protein and scRNA-scATAC integration

To benchmark CHAMPOLLION against current state-of-the-art methods under larger, more realistic conditions and to study how bridge size influences performance, we turned to two gold-standard datasets from the NeurIPS 2021 Open Problems (OP) in single-cell analysis challenge (Figure 2e): (i) OP Multiome, a human bone-marrow dataset containing 69,249 RNA + ATAC profiles collected across 13 site-and-donor batches [34]; and (ii) OP Cite, a human bone-marrow dataset comprising 90,261 RNA + Antibody Derived Tags (ADT) profiles spanning 12 site-and-donor batches [34]. Both datasets are widely regarded as gold standards for multi-omic integration, as they contain ground-truth multimodal pairs and high quality cell type annotations, routinely serving as reference benchmarks for leading methods. Importantly, in real applications, paired and unpaired data are produced by different technologies, often in separate labs and from distinct donors, so a pronounced batch effect between the bridge cells and their unimodal counterparts is unavoidable. The OP datasets enable us to emulate this scenario: we deliberately draw the bridge cells and the pseudo-unpaired cells from non-overlapping batches. To probe how bridge size affects performance, we first divided each dataset roughly 50:50, treating one half as a test set of pseudo-unpaired cells and the other as the bridge. From the latter we drew progressively larger subsamples, ranging from 500 to 33,460 cells for OP Multiome and from 500 to 45,607 cells for OP Cite. This resulted in bridges of increasing sizes, with the smallest setting posing a particularly demanding challenge. The exact splitting and sampling strategy is detailed in the Methods section. Performance was then assessed on the full test set with the same LTA and FOSCTTM metrics described above.

Across both datasets, CHAMPOLLION outperforms the state-of-the-art baselines on both metrics, as shown in Figure 2f. On OP Multiome, MIDAS overtakes Seurat, whereas the ordering reverses on OP Cite; yet CHAMPOLLION always obtains the best scores, including when the full bridge is available. What matters most, however, is performance when paired data are scarce. With only 2,000 bridge cells CHAMPOLLION already holds a considerable lead over the baselines. Remarkably, CHAMPOLLION reaches near-maximal accuracy with only a few thousand bridge cells, after which adding more pairs yields little extra gain. In an even more extreme regime, with only a few hundred paired cells, the LTA margin over baselines narrows thus showing that there is an inherent lower bound for a successful integration. Nonetheless, the FOSCTTM still shows CHAMPOLLION making better alignments, demonstrating that it extracts the maximum value from the limited supervision.

CHAMPOLLION’s strong performance with limited bridge data likely stems from three factors: the integration of informative priors, a simple linear cost model, and, above all, its weighted cost regularization. To illustrate the role of *γ*, we measured performance at several *γ* values for all bridges, as shown in Supplementary Figure 1. With only a few paired cells, setting *γ* too low weakens the prior’s influence and hurts accuracy, whereas *γ* = 0.1 strikes an effective balance. Conversely, when many pairs are available, an excessively high *γ* over-regularizes the problem, preventing the model from fully exploiting the abundant pairing signal and thus lowering performance relative to milder regularization. Users can therefore adjust *γ* according to their expectations about bridge quality and size.

### 2.4 CHAMPOLLION reveals biologically meaningful cross-modal relationships in scRNA-protein data

The benchmark demonstrated CHAMPOLLION’s ability to align unpaired cells across modalities through the transport plan. Moreover, this matching is driven by a learned cross-modal cost whose parametrization provides a feature interaction matrix that captures molecular relationships between modalities. Unlike existing bridge-integration approaches, which do not offer an interpretable feature-level component, our learnt model can be directly investigated to reveal cross-modal links.

In previous experiments, the model was trained on dimension-reduced representations, which prevented direct interpretation of these interactions. In scRNA-scATAC integration in particular, the extreme dimensionality made working directly with features difficult. In CITE-seq data, however, this is possible due to the limited number of measured proteins, thus offering a tractable setting to investigate links between surface proteins and transcripts measured in the same cells (see Figure 3a). To this end, we focus on a single batch of the previously described OP Cite dataset, comprising 10,966 cells. We use 4,031 highly variable genes to represent the RNA modality and the 134 measured surface proteins to represent the ADT modality.

**Fig. 3:**
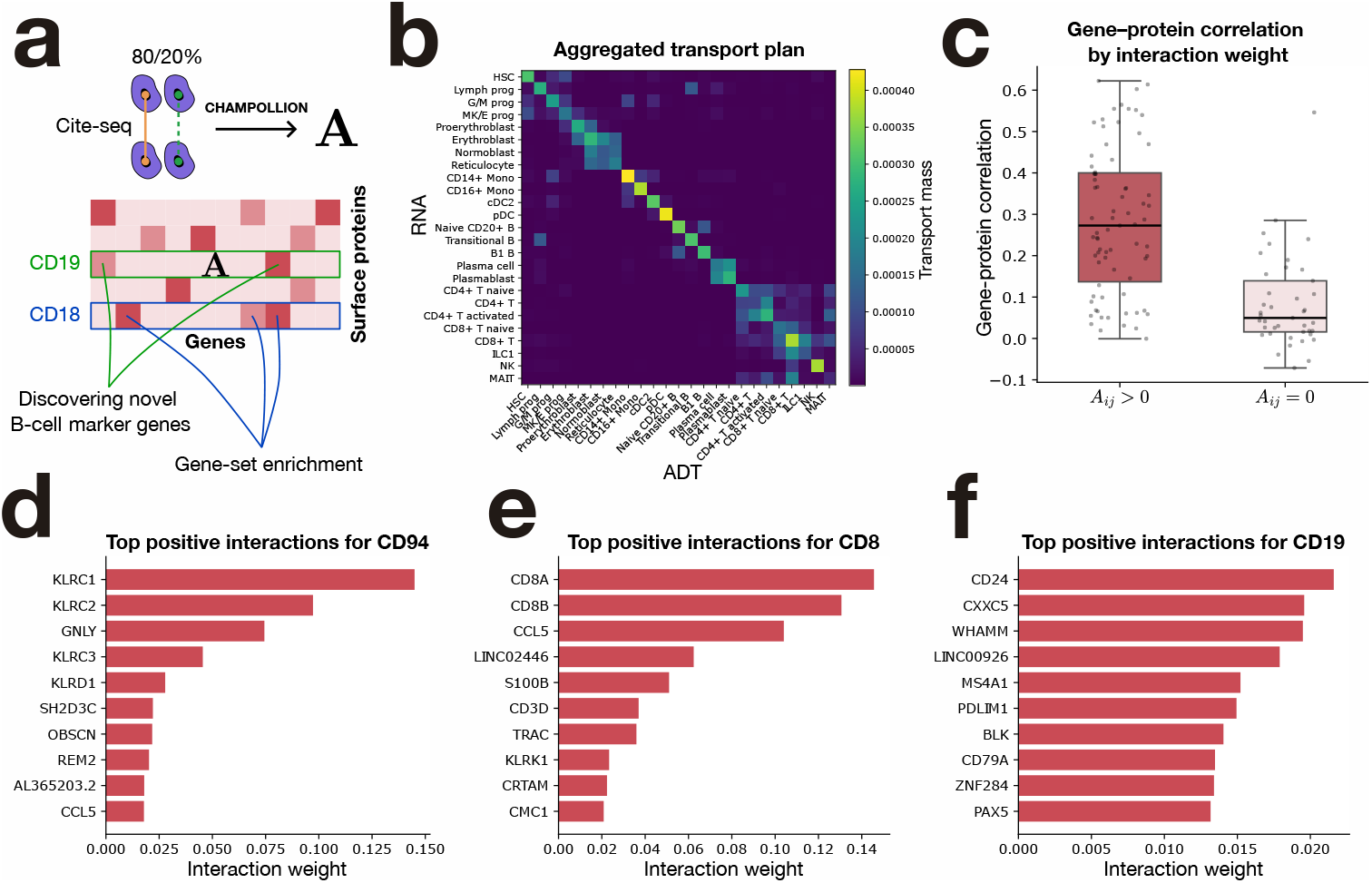
Feature-level integration and interpretability of scRNA-protein data. **(a)** Overview of the integration and interpretation framework. **(b)** Transport plan inferred by CHAMPOLLION on the 20% held-out pseudo-unpaired cells (rows correspond to RNA cells and columns to ADT cells), subsequently sum-aggregated by cell type. **(c)** Coding gene-protein expression correlation for pairs with zero inferred interaction (*n* = 41) versus positive interaction (*n* = 75). **(d-f)** Ten highest positively associated genes with CD94 (d), CD8 (e) and CD19 (f) in the inferred **A** matrix.

To demonstrate first that CHAMPOLLION can integrate cells using feature-level data, we treated 80% of the cells as paired and the remaining 20% as unpaired. As shown in Figure 3b, CHAMPOLLION maintains strong integration performance, achieving an LTA of 0.874 and a FOSCTTM of 0.033 on pseudo-unpaired cells, proving that it can be applied directly to feature-level data while maintaining a good alignment quality.

#### 2.4.1 Learned interactions recover known gene-protein pairs

We now turn to the learned interaction matrix **A** which captures relationships between features whose expression levels are linked within the same cells, potentially for a variety of regulatory or mechanistic reasons. Because such cross-modal dependencies can arise through multiple biological processes, there is generally no clear ground truth against which to evaluate the learned interactions. One class of interactions for which partial prior knowledge exists is coding gene-protein pairs. In this dataset, 116 of the 134 measured surface proteins have their corresponding coding genes represented in the RNA modality. In the benchmark experiments, this correspondence was incorporated as prior information, reflecting the expectation that the expression of a gene is associated with the abundance of its encoded protein. Here, however, to assess what CHAMPOLLION can recover directly from the paired data alone, we ran the method without including this prior cost and studied specifically the entries of the learned matrix **A** corresponding to coding gene-protein pairs. We first observe that all 117 coding gene-protein interactions have non-negative weights in the learned matrix **A**, consistent with the expectation that gene expression should not be anti-correlated with abundance of its encoded protein. Among them, 75 pairs display strictly positive interaction weights, in agreement with the standard prior assumption. The remaining 41 pairs receive a weight of zero. Importantly, this does not indicate an error of the method. It is well established that mRNA levels are not always predictive of surface protein abundance, due to mechanisms such as post-translational regulation, differential trafficking, or variable protein turnover [35]. For example, we find *A*_*ij*_ = 0 for the pair CD152 (CTLA-4). This is biologically coherent, as surface expression of CTLA-4 is known to be tightly regulated through restricted trafficking to the plasma membrane and rapid internalization, such that transcription of the gene does not necessarily result in detectable surface protein levels [36]. More systematically, we examined the 41 gene-protein pairs with *A*_*ij*_ = 0 and found that, in the paired measurements, the correlation between gene expression and protein abundance is significantly lower for these pairs than for those with *A*_*ij*_ *>* 0 (see Figure 3c). This indicates that the learned interactions do not merely reproduce prior assumptions, but instead selectively retain gene-protein links supported by the data while filtering those for which the assumed relationship is not reflected in the measurements.

#### 2.4.2 Cross-modal interactions reveal coherent cell type marker programs

We next sought to demonstrate how CHAMPOLLION can use established markers in one modality to uncover novel markers in the other modality. To do so, we selected surface proteins known to define canonical immune populations and inspected, for each of them, the ten genes with the highest positive interaction weights in **A**. The rationale was that these genes should tend to be co-expressed in the same cells, and therefore may include established markers of the corresponding cell type, as well as potentially less-characterized genes associated with the same population. For CD94, a canonical Natural Killer (NK) cell surface receptor, the top interacting genes reflect well-established NK cell biology (see Figure 3d). We recover KLRD1, which encodes CD94 itself, as well as KLRC1, KLRC2, and KLRC3. These genes belong to the NKG2 family and form heterodimeric receptors with CD94: KLRC1 is associated with inhibitory signaling, whereas KLRC2 and KLRC3 participate in activating receptor complexes [37, 38]. We also identify GNLY (Granulysin), an antimicrobial peptide stored in cytotoxic granules and specifically expressed in cytotoxic lymphocytes such as NK and CD8 T cells [39]. A similar observation can be made for CD8, as shown in Figure 3e. Among the top ten interacting genes, we find nine genes consistent with CD8 T cell identity and cytotoxic function. These include CD8A and CD8B, the transcript-level markers encoding the CD8 co-receptor [40]; CD3D and TRAC, core components of the TCR/CD3 complex and canonical pan-T-cell markers [41]; CCL5, a well-established effector chemokine associated with cytotoxic and memory CD8 T-cell states [42, 43]; KLRK1 (NKG2D), a receptor strongly linked to cytotoxic T and NK functional programs [44]; and CRTAM, an activation-induced receptor described in activated CD8 T cells [45]. We also identify S100B [46] and LINC02446 [47], which have been reported in specific CD8 T-cell subsets rather than representing core CD8 identity, suggesting that the learned interactions may capture more specialized or activation-associated CD8 states in addition to canonical lineage markers. For CD19, a canonical marker of B cells (see Figure 3f), we retrieve five well-established B-lineage markers among the top interacting genes: CD79A, a core component of the B-cell receptor complex expressed early in B-cell development [48]; PAX5, a central transcription factor (TF) maintaining B-cell identity [49]; BLK, a tyrosine kinase with B-lineage-restricted expression [50]; MS4A1 (CD20), a marker of mature non-plasma B cells [51]; and CD24, a molecule associated with B-cell development and naïve B-cell subsets [52]. In addition, we identify LINC00926, a less-characterized non-coding RNA that has interestingly been proposed as a biomarker of naïve B cells in a recent study [53]. Across these examples, the learned interactions recover both lineage-defining markers (such as CD79A and PAX5 for B cells, or CD8A/CD8B for CD8 T cells) and genes associated with functional or effector states (such as GNLY in cytotoxic lymphocytes). The fact that CHAMPOLLION retrieves well-established markers directly from cross-modal interactions validates the biological relevance of the learned matrix. More broadly, this suggests that the method can be used to discover novel or less-characterized markers in one modality by leveraging known markers in another, an especially valuable property in modalities that are more difficult to interpret than transcriptomics such as Raman spectroscopy [54].

#### 2.4.3 Functional programs emerge from surface protein-gene interactions

To demonstrate CHAMPOLLION’s ability to provide functional insights into less-characterized surface proteins, we analyzed the strongest interacting genes for selected proteins and tested whether these gene sets converged on interpretable biological pathways (see Methods).

As a first example, we considered CD36, a broadly expressed surface protein not restricted to a specific lineage. Over-representation analysis of its top interacting genes revealed significant enrichment (*p <* 0.05) for the terms “scavenger receptor activity” and “apoptotic cell clearance” (see Supplementary Table 1). These enrichments are consistent with the known biology of CD36, which belongs to the class B scavenger receptor family and plays a key role in the recognition and phagocytic clearance of apoptotic cells (efferocytosis) [55, 56]. Supporting this interpretation, FCN1, a gene associated with phagocytic processes, appears among the top inferred interactions associated with CD36 [57] (see Supplementary Table 2).

We next investigated CD162 (PSGL-1) who acts as a counter-receptor for E-selectin and mediates leukocyte trafficking to sites of infection or inflammation. Over-representation analysis of its top interacting genes revealed significant enrichment for the terms “cell killing” and “leukocyte mediated cytotoxicity” (see Supplementary Table 1), in line with the inflammatory contexts in which CD162-mediated trafficking occurs. This enrichment is supported by the two strongest inferred interactions (Supplementary Table 2), GNLY and GZMB, two key cytotoxic effector molecules expressed by NK cells and cytotoxic T lymphocytes. Beyond its established role as a homing and adhesion molecule, CD162 has also been studied in the specific context of acute myeloid leukemia (AML), where it has been implicated in the promotion of chemoresistance [58, 59]. In accordance with this, we identify TNFAIP3 (A20) whose elevated expression has been associated with induction failure and anthracycline resistance in AML [60]. We also recover PIM1, a kinase reported to contribute to cytarabine resistance in AML models [61]. Taken together, these findings underline the therapeutic relevance of CD162 in AML and point toward a coordinated adhesion-survival signaling axis.

As a final example, we examined CD18 (encoded by ITGB2)a *β*2 integrin subunit that plays a central role in immune cell adhesion and activation. We identify CD44 among top genes associated with CD18 (see Supplementary Table 2), which echoes evidence that crosslinking CD44 can induce activation of ITGAL/ITGB2 complexes [62]. Over-representation analysis of the top interacting genes further revealed significant enrichment for the “Senescence and autophagy in cancer” pathway (see Supplementary Table 1). In line with this result, a recent study suggests that ITGB2 may regulate mTOR expression through the PI3K-AKT pathway, thereby promoting cellular energy supply, and inhibiting mitophagy in ovarian cancer cells [63]. Importantly, ITGB2 has been implicated in multiple cancer contexts beyond ovarian cancer, including triple-negative breast cancer [64] and melanoma [65], where experimental evidence links it to tumor progression. This thus raises the possibility that adhesion-dependent metabolic reprogramming mediated by ITGB2 may represent a broader mechanism active across tumor types.

Altogether, these examples illustrate that the learned interaction matrix captures biologically and clinically relevant surface protein-gene associations, highlighting CHAMPOLLION’s ability to uncover meaningful cross-modal relationships beyond canonical lineage markers.

### 2.5 CHAMPOLLION integrates gene expression and chromatin accessibility at atlas scale in human tonsils

The Human Cell Atlas initiative [66] has driven the generation of large-scale single-cell atlases aimed at systematically mapping cellular diversity across human tissues. While early atlases primarily relied on single-cell RNA sequencing to produce transcriptional maps of tissues, more recent efforts incorporate complementary modalities such as chromatin accessibility (scATAC-seq), providing regulatory insights. The increasing availability of extremely large single-cell atlases spanning multiple molecular layers further highlights the need for multimodal integration methods. Their scale offers unprecedented resolution and statistical power to refine cellular identities, yet renders many existing computational methods impractical for datasets comprising hundreds of thousands to millions of cells.

The Human Tonsil Atlas (HTA) [67] represents a particularly rich example of such efforts, where in addition to standalone scRNA-seq and scATAC-seq datasets, it includes a multiome dataset with paired RNA and ATAC measurements in the same cells. This configuration provides an ideal large-scale testbed for CHAMPOLLION, which leverages paired measurements to learn a cross-modal metric that can subse-quently be deployed to integrate the larger unpaired cohorts, enabling a coordinated dissection of immune cell states through combined transcriptional and regulatory signals (see Figure 4a). In the data analyzed here, the multiome measurements comprised 68,749 paired RNA-ATAC cells, alongside 209,774 scRNA-seq cells and 58,049 scATAC-seq cells. As in our benchmarking setting, the extreme dimensionality of the data, most notably the 215,000 ATAC peaks, combined with its scale, motivated the use of low-dimensional embeddings derived from DRVI [68] as inputs to CHAMPOL-LION. We embedded RNA and ATAC profiles using separate DRVI models and, with these representations, were able to learn the cross-modal cost and integrate the full dataset in under 12 hours.

**Fig. 4:**
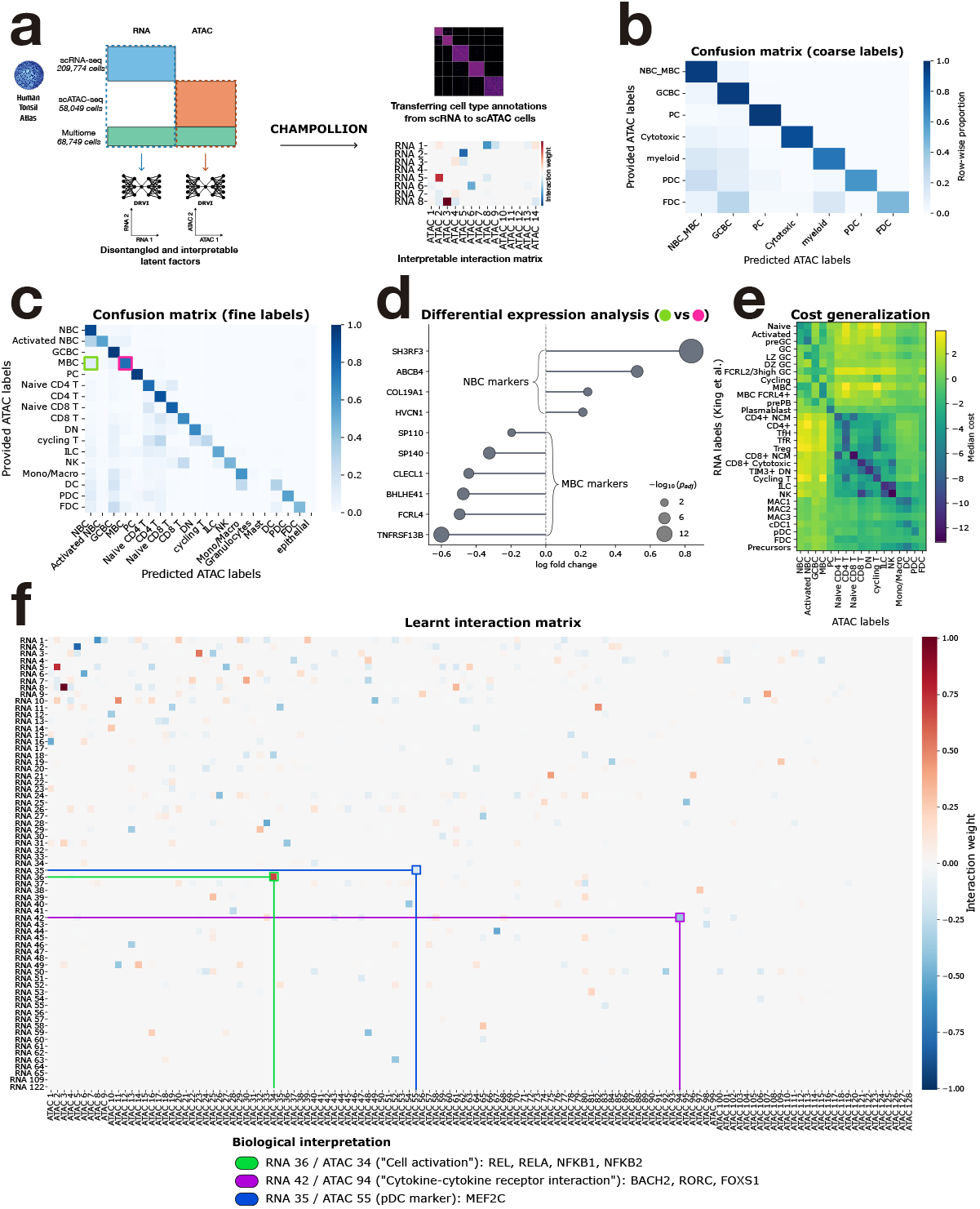
Atlas-scale integration of RNA and ATAC data in human tonsils. **(a)** Schematic representation of the integration. **(b)** Row-wise normalized confusion matrix comparing coarse cell type labels transferred from RNA to ATAC cells by authors of the dataset (rows) with labels inferred by CHAMPOLLION (columns). **(c)** Same as (b) using finer annotations. Highlighted groups correspond to cells labeled as memory B cells by the HTA authors and reassigned by CHAMPOLLION to memory B cells (pink, 8401 cells) or naive B cells (green, 1262 cells). **(d)** Differential expression analysis on gene activities between the highlighted groups in (c), showing selected naive and memory B cell marker genes. The x-axis indicates log fold change and point size indicates adjusted *p*-values obtained with a t-test (see Methods). **(e)** Cross-modal cost between HTA ATAC cells (columns) and an external human tonsil scRNA-seq dataset (rows), computed using the inferred **A** and aggregated by cell type (median), illustrating generalization of the learned cost. **(f)** Learned interaction matrix **A** between RNA and ATAC latent factors (rows and columns, respectively). Highlighted pairs of latent factors are annotated with enriched pathways and TFs.

We first assessed the ability of CHAMPOLLION to align cells across modalities. Because the scRNA-seq and scATAC-seq datasets are unpaired, direct cell-to-cell validation is not possible. Instead, we evaluated integration quality using provided cell type annotations. In this atlas, cell identities were manually annotated in the scRNA-seq and multiome RNA data and subsequently transferred computationally to the scATAC-seq dataset. As ATAC labels are therefore inferred rather than ground truth, the LTA should be interpreted as a measure of concordance with the atlas annotation procedure rather than absolute correctness. The atlas provides annotations at multiple levels of granularity, with 9 coarse cell classes and 23 finer subtypes, enabling us to evaluate concordance between methods at different resolutions.

As shown in Figure 4b, the LTA of 90.9% indicates that at the level of coarse cell classes CHAMPOLLION yields predictions largely consistent with the atlas annotation procedure (see Supplementary Figure 2 for the confusion matrix with counts). At the finer resolution, concordance remained high with an LTA of 85.5%, although lower than with coarse annotations, consistent with the increased complexity of transferring annotations among closely related immune subtypes. We next examined the largest source of disagreement for fine cell types (see Figure 4c and Supplementary Figure 3 for the confusion matrix with counts), which involved 1,262 cells annotated in the atlas as memory B cells (MBC) but predicted by CHAMPOLLION as naïve B cells (NBC). To investigate this discrepancy, we performed differential expression analysis on log-normalized gene activities between these 1,262 cells and the subset of 8,401 cells consistently annotated as MBC by both approaches, as shown in Figure 4d. The disputed cells displayed over-expression of established naïve B cell markers, including SH3RF3, ABCB4, HVCN1 and COL19A1, together with under-expression of canonical memory B cell markers such as FCRL4 and TNFRSF13B [67, 69]. These results suggest that the observed disagreement does not reflect misclassification, but may instead highlight cases in which CHAMPOLLION refines annotation transfer between closely related B cell states.

To further assess whether the learned cross modal metric captures biologically meaningful structure, we compared scATAC cells, annotated using CHAMPOLLION, to an independent scRNA-seq dataset that had been annotated separately [70]. This external reference provides an orthogonal measurement, enabling us to evaluate whether costs induced by the learned metric are coherent beyond the original atlas framework. Although annotation categories do not map perfectly one to one across datasets, the cell type-aggregated cross-modal cost shown in Figure 4e reveals lower costs between corresponding cell types across datasets. More precisely, for 23 out of 29 RNA cell types (excluding “Precursors” due to the ambiguous annotation), the closest ATAC cell type under the learned metric corresponds to a biologically consistent match, based on a set of allowed correspondences between cell type annotations (see Supplementary Table 3). For instance, CD4+ NCM (naive/central memory) cells in the scRNA-seq dataset are closest to T lymphocyte populations in the ATAC data, with increasing proximity to more specific subsets, and ultimately exhibit the lowest cost to naïve CD4 T cells. Together, these results indicate that the learned metric captures biologically coherent cell similarities and generalizes beyond the original annotation transfer setting.

The interpretability of each modality’s DRVI latent space then enabled us to interpret the learned cross-modal relationships. Indeed, DRVI models produce latent factors that capture independent biological signals, each of which can be interpreted through the molecular features most strongly associated with it. Because CHAMPOL-LION learns interactions between RNA and ATAC latent factors through the matrix **A** (Figure 4f), the resulting cross-modal cost can be interpreted as relationships across modalities between biological processes. Associating RNA and ATAC latent factors through their strongest interaction in **A** revealed biologically coherent connections between the processes captured in the two modalities. In some cases, both latent factors were highly specific to a particular cell population, for example RNA 21 and ATAC 75 are both predominantly active in naïve CD8 T cells (see Supplementary Figures 4 and 5 for visualizations of the latent factors’ activities per cell type). In other cases, the associated latent factors captured broader biological programs shared across multiple cell types, such as RNA 13 and ATAC 18 which are associated with cell cycle activity (see Supplementary Table 4).

To further characterize these relationships, we investigated several highly interacting pairs of latent factors. For each pair, we analyzed the genes associated with the RNA latent factor and evaluated pathway enrichment among these genes. Additionally we identified TFs associated with each latent factor separately, based on genes linked to the RNA factor and on TF motifs enriched in ATAC peaks associated with the paired ATAC factor, and focused on the TFs shared between the two sets (see Methods). This analysis allowed us to assess whether the learned interactions correspond to coherent regulatory programs across modalities.

One illustrative example involves the latent factor RNA 36, which shows its strongest interaction with ATAC 34, both being specifically active in the activated naïve B cell cluster. Genes associated with RNA 36 are enriched for the Gene Ontology term “cell activation” (see Supplementary Table 4). Moreover, the intersection of TFs enriched in the RNA and ATAC latent factors is composed of REL, RELA, NFKB1 and NFKB2 (see Supplementary Table 5), key components of the NF-kB pathway known to play central roles in B cell activation [71, 72].

A second example involves RNA 42 and ATAC 94. In contrast to the previous case, these latent factors are active across multiple immune populations, including innate lymphoid cells and T cells, indicating a broader program. Genes associated with RNA 42 are enriched notably for the KEGG pathway “Cytokine-cytokine receptor interaction” (see Supplementary Table 4). We identified BACH2 and RORC among the TFs enriched in both modalities, as shown in Supplementary Table 5. This observation is consistent with the role of BACH2 in regulating cytokine signaling responses across T, B and innate lymphoid cells [73–75], while RORC is a master TF best known for controlling Th17 and ILC3 differentiation downstream of cytokine signaling pathways such as IL-23 [76].

Finally, we focused on RNA 35 and ATAC 55 which are both highly specific to plasmacytoid dendritic cells, as shown in Supplementary Figures 4 and 5. Although no pathway enrichment reached significance for the genes associated to RNA 35, the intersection of TFs enriched in both modalities contained a single candidate, MEF2C (see Supplementary Table 5). This finding is notable given previous work showing that MEF2C can restrain inflammatory activation by limiting NF-*κ*B nuclear translocation in microglia [77] and protect against the development of atherosclerosis by inhibiting TLR/NF-*κ*B signaling in endothelial cells [78]. As NF-*κ*B signaling is also central to plasmacytoid dendritic cell activation and cytokine production [79, 80], this observation raises the possibility that MEF2C may play a broader role in regulating inflammatory responses beyond the brain, including in plasmacytoid dendritic cells as suggested by our integration.

## 3 Discussion

Anchoring large single-omic datasets with a modest multi-omic bridge, also known as bridge integration, offers a practical route to deep, cross-modal single-cell analysis for most laboratories. Yet current methods typically couple modalities solely through the bridge, often ignoring feature-level priors, and lack explicit regularizing mechanisms to control the influence of the paired data, making them sensitive to biases and incomplete coverage. In addition, they offer limited interpretability of the inferred cross-modal relationships.

CHAMPOLLION fills this gap by casting bridge integration as an inverse optimal transport problem. It learns an interpretable cross-modal metric from the paired cells, complemented by feature-level priors, and then uses that metric to align the remaining unimodal cells. A tunable regularization term balances the weight of the bridge against the priors, giving CHAMPOLLION resilience to bridges that are small or incomplete.

Through rigorous benchmarks on gold-standard RNA+ATAC and RNA+ADT datasets, we showed that CHAMPOLLION outperforms state-of-the-art alternatives. Crucially, it stays accurate with tiny or cell type limited bridges, requiring only a few thousand paired cells to match the performance it achieves with bridges tens of thousands strong, while its tunable regularization ensures that abundant bridges are fully exploited when available. We further illustrated the biological relevance of CHAM-POLLION through two case studies. In RNA-protein data, the learned cross-modal cost recapitulated meaningful gene-protein relationships, recovering known pairs while also enabling discovery of novel cell type markers and functional insights for less-characterized proteins, including a potential role for CD18 in tumor progression across multiple cancers. In a RNA-ATAC human tonsil atlas, CHAMPOLLION successfully integrated large datasets and provided mechanistic insights, notably suggesting that MEF2C may act as a broader regulator of inflammatory responses beyond the brain, including in plasmacytoid dendritic cells.

The transport plan inferred by CHAMPOLLION supports downstream analyses such as annotation transfer without requiring a shared latent embedding. However, as it relies on classical optimal transport, it assumes balanced distributions between modalities. In practice, differences in cell type composition arising from sampling biases, sorting strategies, or biological conditions can violate this assumption and distort correspondences due to the mass-preservation constraint. Notably, this limitation affects only the matching step, while the learned cross-modal cost, estimated from paired and thus balanced data, remains unaffected and can be used independently without inferring a transport plan, for instance for annotation transfer. A more principled solution nonetheless would be to incorporate unbalanced optimal transport (uOT), which relaxes the mass constraint and enables robust integration in the presence of imbalanced or missing cell populations [27, 28, 81].

In addition, CHAMPOLLION is currently restricted to the integration of two modalities, whereas recent methods such as MIDAS address the more general setting of mosaic integration, defined as the integration of arbitrary combinations of paired and unpaired datasets across multiple modalities. Although CHAMPOLLION is not designed for this general setting, extending it to handle more than two modalities would be particularly relevant given emerging technologies such as DOGMA-seq or TEA-seq [6, 82], which profile three modalities in the same cells. While optimal transport naturally extends to multi-marginal formulations [83], their computational complexity increases rapidly with the number of modalities and would require methodological adaptations to remain tractable. Exploring such extensions represents an exciting direction for future work.

A key strength of CHAMPOLLION lies in its modular design, which allows it to operate on arbitrary single-cell representations. In this work, we demonstrated its effectiveness using both raw features and learned embeddings, suggesting that it can readily benefit from ongoing advances in single-cell representation learning [84, 85]. More broadly, this modularity extends beyond the choice of representation to the modalities themselves. By learning cross-modal relationships, CHAMPOLLION enables the transfer of information across modalities, both at the cell level (e.g., annotation transfer) and at the feature level, where it can uncover functional links or suggest novel markers in one modality based on signals from another. This flexibility makes CHAMPOLLION particularly well suited to emerging multimodal technologies that couple transcriptomics with more complex and less directly interpretable readouts. Recent developments such as RamanOmics [54], which combines Raman spectroscopy with single-nucleus RNA-seq, provide access to biochemical and biophysical properties of cells at the level of molecular bonds. This includes lipid composition, protein structure and metabolic states, features that are largely invisible to genomic assays yet critically shape cellular phenotypes. Similarly, approaches such as IRIS [86] integrate high-resolution imaging with single-cell transcriptomics, capturing rich morphological information about cellular architecture with direct functional and clinical relevance. In both cases, the complementarity between modalities offers a unique opportunity to link gene expression programs to biochemical or morphological phenotypes. Methods such as CHAMPOLLION, which can leverage these complementary signals while providing interpretable feature-level relationships, therefore hold strong potential for uncovering new principles of cellular identity in these next-generation multimodal datasets.

## 4 Methods

### 4.1 Optimal Transport

The classical Optimal Transport (OT) problem, defined by Monge [87] and Kantorovitch [88], aims at comparing two probability distributions by finding a transport plan that moves one distribution onto the other at minimal cost. In our setting, both distributions are discrete measures, written 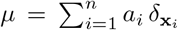 and 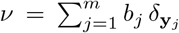, where 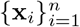 and 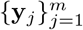 denote the samples and 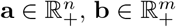 are weights such that ∑_*i*_ *a*_*i*_ = ∑_*j*_ *b*_*j*_ = 1 (which in our study will always be uniform, i.e. 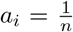 and 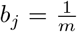). This cost is specified by a function *c* measuring the effort required to move mass between individual points **x** and **y**. We denote by *C* ∈ ℝ^*n×m*^ the cost matrix defined entrywise as *C*_*i,j*_ = *c*(**x**_*i*_, **y**_*j*_).

The OT problem formally reads as:

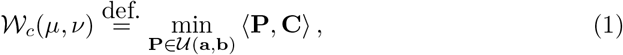

where 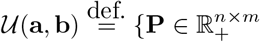 such that ∑_*j*_*P*_*i,j*_ = *a*_*i*_ and ∑_*i*_*P*_*i,j*_ = *b*_*j*_}. The transport plan **P** encodes how mass is transported across the two distributions, with *P*_*ij*_ representing the amount of mass moved from **x**_*i*_ to **y**_*j*_, and the objective ⟨**P, C**⟩ corresponding to the total transport cost.

Adding an entropic regularization weighted by a parameter *ε* to the objective function of Equation 1 results in a new optimization problem noted:

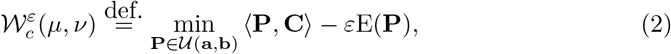

where E: **P** 1→ − ⟨**P**, log **P** − **1**⟩ is the Shannon entropy.

While setting *ε* = 0 recovers the unregularized OT problem (Eq. 2), using *ε >* 0 makes the problem *ε*-strongly convex. It can be solved computationally much faster than its unregularized counterpart with the GPU-enabled Sinkhorn algorithm [89].

This problem admits a dual formulation, defined as follows.

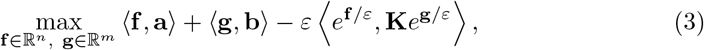

where 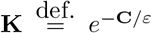 (the exponential is applied element-wise) is the Gibbs kernel associated with the cost function *c*.

Furthermore, by leveraging the closed-form expression for the optimal **g**, given **f** one can obtain the following semi-dual formulation:

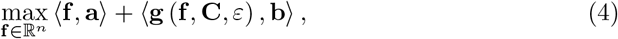

where 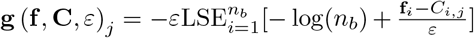 with LSE referring to the log-sum-exp operation 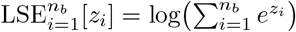.

### 4.2 Inverse Optimal Transport

A crucial aspect of OT is the cost function *c*, in standard OT such as Equation 1 or Equation 2, *c* is known and leveraged to provide a soft matching **P** of the samples. Inverse Optimal Transport aims to answer the opposite question: given a matching between samples, can we learn a cost function *c* which explains this matching [22]?

This is formalized as a bi-level optimization problem, This is formalized as a bi-level optimization problem, where the objective depends on the coupling **P**_*c*_, which is itself defined as the solution of Equation 2 parameterized by *c*. We then estimate *c* by minimizing the Kullback–Leibler divergence KL between an observed coupling 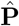 and the induced coupling **P**_*c*_, optionally adding a regularization term ℛ (*c*):

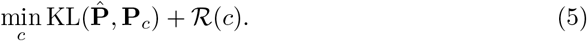

As shown in [22], this minimization problem is convex with respect to *c*. Equivalently, this problem can be read as:

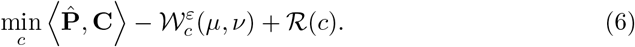

By then replacing the entropic OT term with it’s semi-dual formulation Equation 4, we obtain a single-level and more tractable problem:

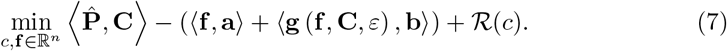

### 4.3 CHAMPOLLION

CHAMPOLLION takes as input paired and unpaired single-cell data from two modalities, labeled 1 and 2. The paired measurements or “bridge” are contained in count matrices 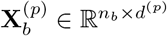 with *p* = 1, 2, measured on the same *n*_*b*_ cells, while the larger unimodal measurements to integrate are 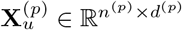, measured on unpaired sets of cells. We use the notation 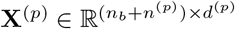 to describe the full dataset obtained by concatenating bridge and unpaired cells along the cell dimension, 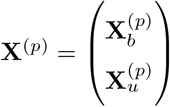. Leveraging feature-level priors, we also have access for both paired and unpaired cells to additional measurements expressed in a feature space common to both modalities 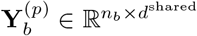 and 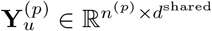, which provides a common basis for comparing cells across modalities. CHAMPOLLION first uses inverse optimal transport (iOT) to learn a cost function from the paired measurements and then employs this cost function to match unpaired cells with optimal transport.

In more details, the goal of the first stage of the method is to learn the following parametric cost function: 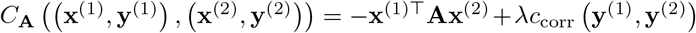 where **x**^(*p*)^ (respectively **y**^(*p*)^) is a single cell represented by a row of the complete **X**^(*p*)^ (respectively **Y**^(*p*)^) matrix, *c*_corr_ is a correlation-based dissimilarity, and the matrix **A** captures feature associations across modalities which we want to optimize, with the negative sign ensuring that positive entries in **A** correspond to positive associations between features. To estimate **A**, we regard the bridge as a known transport plan, corresponding to the empirical coupling 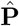 introduced above. Since paired cells are aligned in the same order in 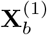 and 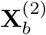, this coupling is given by the normalized identity matrix 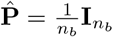, and we solve the associated iOT problem (Equation 7). We denote by 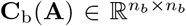 the cost matrix defined by 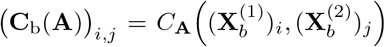 i.e. obtained by evaluating *C*_**A**_ on all cross-modal pairs within the bridge. Similarly, we define 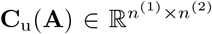 as 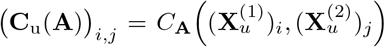 the corresponding matrix for the unpaired cells.

We thus solve:

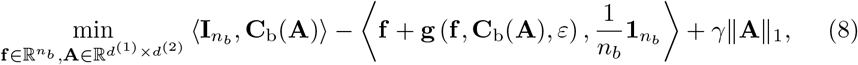

where we choose an *ℓ*_1_ penalty to regularize the learnt cost: ℛ (*C*_**A**_) = *γ*∥**A**∥_1_. This formulation offers a convex objective which can be optimized efficiently with standard first-order optimization methods.

Once the parameter **A** is fitted, we can obtain a transport plan **P** that matches unpaired cells by solving a classical entropic optimal transport problem (Equation 2) with **C** = **C**_u_(**A**) as the ground cost.

### 4.4 Implementation details

A difficulty in optimizing Equation 8 is that the *ℓ*_1_ penalty is not differentiable, which typically requires the use of an iterative soft-thresholding algorithm [24]. To have a more scalable solver, we follow [25] and resort to an over-parameterization method leading to a smooth optimization problem. Indeed, Lemma 1 of [90] shows that

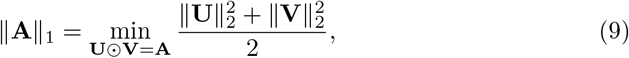

where ⊙ denotes the element-wise product. This allows us to reparameterize any problem of the form min_**A**_ *H*(**A**) + *γ*∥**A**∥_1_ as 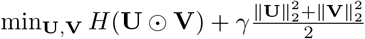. This formulation is smooth and can be optimized with gradient-based methods, after which **A** is recovered as **U** ⊙ **V**.

We implement CHAMPOLLION in PyTorch [91] and, optionally for large datasets, wrap pair-wise computations in PyKeOps LazyTensors [29], which keep large matrices symbolic and generate their values on-the-fly during GPU-accelerated reductions, greatly lowering memory use.

To solve Equation 8, we use Torch’s `Adam` Optimizer [92] with default parameters and a learning rate of 0.001. Iterations are stopped once the convergence criterion is met, which is achieved when the *ℓ*_1_-norm of the gradient with respect to **f** of the objective function in Equation 8 is lower than 0.001.

For the standard entropic OT problem in the second stage, we re-implemented the Sinkhorn algorithm.

In all experiments, we used the following default parameters: *γ* = 0.01, *ε* = 1, *λ* = 20. We studied the impact of the choice of *γ* on performance depending on different data regimes in Supplementary Figure 1. However we set *ε* = 1 by default and did not explore any other values, as the objective function in Equation 8 is 1-homogeneous, i.e., scaling *ε* is equivalent to rescaling (**A, f**). For the choice of *λ*, note that it controls the relative weight of the prior cost. In the over-regularized regime (large *γ*), where **A** = 0, the resulting cost reduces to *λ* times the prior cost. We choose *λ* so that this quantity has an appropriate scale for *ε* = 1. Since the cost is mean-normalized and a standard heuristic is to set *ε* ≈ 0.05 × mean(**C**), this amounts to choosing 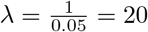.

### 4.5 Unimodal data preprocessing

For preprocessing, we used the Muon [93] and Scanpy [94] packages.

#### 4.5.1 scRNA preprocessing

We performed quality control filtering of cells on the number of expressed genes, and the total number of counts (using Muon’s filter obs). Quality control filtering of genes was performed on the number of cells expressing the gene (using Muon’s filter var). We then applied the log-normalization which consists of normalizing counts for each cell so that they sum to 10,000 (using Scanpy’s normalize total) and then log transforming them (using Scanpy’s log1p). Finally, the log-normalized counts were scaled before being used to compute the first 50 principal components.

#### 4.5.2 scATAC preprocessing

We performed quality control filtering of cells on the number of open peaks and the total number of counts (using Muon’s filter obs). Quality control filtering of peaks was performed on the number of cells where the peak is open (using Muon’s filter var). We then log-normalized the peak counts as described for the scRNA, scaled the lognormalized counts and computed the first 50 principal components.

#### 4.5.3 ADT preprocessing (Cite-seq)

Since the number of measured proteins is small and this data is less noisy than scRNA or scATAC, no quality control or feature selection was performed. We only normalized the data using Muon’s implementation of the Center Log Ratio (CLR) technique.

### 4.6 Computing the prior-knowledge cost

To build the **Y**^(*p*)^ matrices, we first derive shared feature spaces for each modality pair.

For RNA + ATAC, we use `Signac` [95] to compute gene-activity scores from the ATAC profiles, intersect these genes with those measured by scRNA-seq, and retain the top 100 principal-component scores for every cell (computed jointly on RNA counts and gene activities).

For RNA + ADT, we manually map each surface protein to its coding gene using GeneCards, then restrict both modalities to the resulting gene-protein pairs.

The prior cost is then simply the correlation distance (one minus Pearson’s *r*) between every cell in modality 1 and every cell in modality 2, computed with `scipy.cdist` [96].

### 4.7 Splitting datasets between paired and unpaired cells

To assess alignment quality at single-cell resolution, we began with datasets in which every cell is measured in both modalities. To simulate the bridge-integration scenario, we then designated a subset of these paired cells as “unpaired,” leaving the remainder to serve as the bridge. The specific split procedure is detailed below for each dataset.

#### 4.7.1 Human PBMC Multiome dataset

We began with the 10x Genomics Multiome (RNA + ATAC) dataset available at https://www.10xgenomics.com/datasets/pbmc-from-a-healthy-donor-no-cell-sorting-10-k-1-standard-2-0-0 and retained only cells from five populations: memory CD4 T cells, naïve CD8 T cells, classical monocytes, CD56-bright NK cells, and CD56-dim NK cells. For the split, we placed half of the memory CD4 T, naïve CD8 T, and classical monocyte cells into the bridge, while assigning all NK cells and the remaining halves of the CD4 T, CD8 T, and monocyte cells to the unpaired set.

#### 4.7.2 OP Multiome & Cite

For the NeurIPS 2021 Open Problems datasets used in the benchmark, we defined the bridge at the batch (donor × site) level.

- OP Multiome. Batches s4d8, s1d2, s2d1, s3d6, s3d7, s1d3, s2d5, comprising 33 460 cells, served as the bridge; all remaining batches were treated as unpaired.
- OP Cite. Batches s3d7, s3d1, s2d5, s2d2, s1d3, s4d1, s4d8, comprising 45 607 cells, formed the bridge, with the rest considered unpaired.

To assess sensitivity to bridge size, we drew nested random subsamples representing 5 %, 10 %, 20 %, 40 %, 70 %, and 100 % of each full bridge, sampling uniformly across batches with larger subsets containing all cells from smaller ones. As an additional stress test, we created a single-batch bridge by randomly selecting 500 cells from one batch, s4d8 for OP Multiome and s3d7 for OP Cite.

### 4.8 Obtaining shared visualizations with the Transport Plan

Given a transport plan 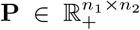 between two modalities, we obtain a shared representation by projecting one modality onto the other using barycentric projection.

For example, the projection of modality 1 onto modality 2 is given by

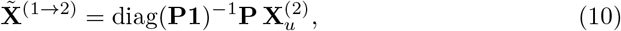

where **1** denotes a vector of ones, so that each projected point is a barycentric combination of samples in modality 2. This yields representations of both modalities in a common space, after which standard dimensionality reduction methods such as UMAP can be applied jointly to all cells as we did in Figure 2c.

### 4.9 Baselines

#### 4.9.1 MIDAS

We compared CHAMPOLLION to MIDAS using the python package *scmidas v0*.*1*.*12*. MIDAS is based on variational architecture, with a modular encoder composed of one neural network per modality. When one cell is profiled in more than one modality, a Product-of-experts module combines modality specific posterior distributions into one latent posterior distribution. Joint posterior regularization and information-theoretic approaches help disentangle biological state from technical noise in latent variables and to align different modalities in the latent space. We followed the official documentation of the package and advice from the authors to run the method on the benchmark datasets.

#### 4.9.2 Seurat

We compared CHAMPOLLION to Seurat’s bridge integration method using the R package *Seurat v5*.*1*.*0*. To integrate unpaired unimodal datasets using a paired bridge, Seurat first performs dictionary learning, representing each unimodal cell as a linear combination of the paired-bridge cells. It then projects these coefficient vectors into a low-dimensional space defined by the leading eigenvectors of the bridge dataset’s cell-cell graph Laplacian. We followed the official documentation to run Seurat on the benchmark datasets.

### 4.10 Evaluation metrics

In our benchmark, we assume that we have *n* cells per modality, that they are paired, and come with one-hot encoded cell type annotations **U** ∈ {0, 1}^*n×c*^ where *c* is the number of cell types. Both the pairing information and the cell type annotations are only used during evaluation to compare the methods’ outputs. We have on one hand embedding methods which output for 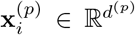, the *i*-th cell in modality *p* (with *p* = 1 or *p* = 2), a low dimensional embedding 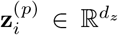. On the other hand, CHAMPOLLION’s evaluated output is a transport plan **P** ∈ ℝ^*n×n*^. Both types of output can be leveraged to obtain a cell-by-cell dissimilarity matrix *D* used as input of the evaluation metrics. For embedding methods, 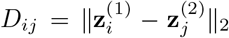, for CHAMPOLLION we can simply use **D** = −**P**.

#### 4.10.1 LTA

The Label Transfer Accuracy (LTA) assesses how effectively cell type labels can be propagated from one modality to the other. We first treat modality *p* as the reference and the other modality *p*^*′*^ = 3 − *p* as the query, predict labels for all query cells, and record the accuracy; we then swap the roles of *p* and *p*^*′*^ and report the average of the two accuracies. For all methods, predictions can use a 5-nearest-neighbours classifier built on the dissimilarity matrix **D**. While this is used for MIDAS and could be used for all methods, CHAMPOLLION and Seurat are treated differently since they both provide direct ways to transfer annotations across modalities.

For Seurat, when RNA is the reference, we employ Seurat’s “TransferData” method, which is more efficient than 5-NN on the embeddings. Seurat’s final score is therefore the mean of (i) 5-NN accuracy when RNA serves as the query and (ii) its built-in transfer accuracy when RNA serves as the reference.

For CHAMPOLLION, the output being a joint distribution coupling cells from both modalities, we can transfer annotations by computing the conditional likelihood of cells belonging to different cell types. To do so, we compute 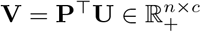. Each row **v**_**j**_ is thus a vector summing to 1 which can be interpreted as the probabilities of cell *j* belonging to each cell type, we thus select the label with the maximal weight. To perform the prediction in the other direction we can repeat the operation using **P** instead of **P**^⊤^.

#### 4.10.2 FOSCTTM

The Fraction Of Samples Closer Than the True Match (FOSCTTM) is a standard metric to evaluate the alignment of multimodal datasets when the ground-truth pairing is known. For each cell *i* in a modality, it quantifies how many cells in the other modality are inferred to be closer to *i* than its true match, this score is then averaged across all cells and also computed in the other direction. More formally, assuming that the *i*-th cell in modality 1 is the true match of the *i*-th cell in modality 2, the FOSCCTM is defined as:

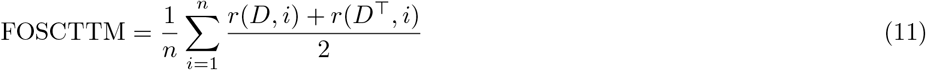

where 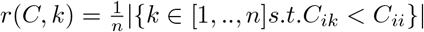

### 4.11 Feature-level interpretability in scRNA-surface protein integration

#### 4.11.1 Preprocessing

For this case study we selected only cells from the largest batch (“s3d6”) of the Cite OP dataset. We randomly selected 80% of the cells as the bridge and used the remaining 20% as the pseudo-unpaired set. For this analysis we didn’t use any prior data and each modality’s counts were preprocessed as in the benchmark (log-normalization for RNA and CLR normalization for ADT) with the addition of a final step: each modality was centered and to unit variance scaled feature-wise, preventing the entries of **A** from being skewed by heterogeneity in feature variances.

#### 4.11.2 Over-representation analysis on a protein’s top gene interactions

For each protein, we identified its most strongly associated genes based on the learned interaction matrix **A**. Specifically, for a given protein, we considered all genes whose absolute interaction score exceeded a fraction of the maximum absolute score for that protein (here, 1*/*100). This relative threshold ensures that selected genes have interaction strengths comparable to the strongest associations, while remaining robust to differences in scale across proteins. To avoid overly large gene sets, we further limited the selection to at most 100 genes, retaining those with highest absolute interaction scores. On the obtained set of genes, we then performed over-representation analysis using the Python interface of the gProfiler service [97], restricting the analysis to the KEGG, Reactome (REAC), WikiPathways (WP), and Gene Ontology (GO) data sources. To avoid bias, we defined the background as the set of 4,031 highly variable genes used for the RNA modality.

### 4.12 Human Tonsil Atlas

#### 4.12.1 Preprocessing

After downloading the Human Tonsil Atlas data using the HCATonsilData package, we retained the discovery cohort for the scRNA-seq data (209,774 cells), alongside 58,049 scATAC-seq cells and 68,749 multiome cells. ATAC profiles were processed with Signac to compute gene activity scores, which were then used to derive the prior cost as described above.

We then used DRVI to obtain low-dimensional embeddings for both modalities, training one model on RNA profiles (scRNA-seq and RNA from multiome) and one on ATAC profiles (scATAC-seq and ATAC from multiome). To enable downstream application of the learned cost to external data, we included 32,607 cells from [70] in the RNA model, ensuring that these cells were embedded in the same space as the HTA data. RNA profiles were restricted to genes shared across datasets, log-normalized (for feature selection only), and reduced to 4,000 highly variable genes using scIB’s hvg batch function to account for intra- and inter-dataset batch effects. The DRVI model was then trained on raw counts with a latent dimension of 128 and hidden size 128 (other parameters left at default).

For ATAC, peak counts were log-normalized (only for feature selection) and 37,561 highly variable peaks were selected using scanpy’s highly variable genes function (min mean=0.01, max mean=2.0, min disp=0.5). Counts were then converted to fragment counts for DRVI, which was trained with a latent dimension of 128, hidden size 256, and a “poisson orig” likelihood as recommended by the authors. In both cases, we used a relatively large latent dimension, as non-informative dimensions are naturally suppressed during training and removed in post-processing.

#### 4.12.2 Differential expression analysis in B cells

To investigate the discrepancy between our transferred annotations and those of the atlas, we performed differential expression analysis on log-normalized gene activities between the 1,262 cells annotated as NBCs by CHAMPOLLION (and as MBCs in the atlas) and the 8,401 cells consistently annotated as MBCs by both approaches. Differential expression was computed using Scanpy’s rank genes groups function with default parameters, which performs a two-sided t-test with Benjamini–Hochberg correction for multiple testing.

#### 4.12.3 Interpretation of the latent factors

For each modality, we used DRVI’s traverse latent and calculate differential vars functions to identify features associated with each latent factor, yielding a score for every feature.

For over-representation analysis on the RNA factors, we selected genes by thresholding these scores using the same relative criterion as in the RNA–protein analysis, retaining genes whose absolute score exceeded 1*/*100 of the maximum for the corresponding factor and limiting the set to the top 100 genes. We then performed over-representation analysis as decribed before using g:Profiler, restricting to KEGG, Reactome (REAC), WikiPathways (WP), and Gene Ontology (GO), with the set of 4,000 highly variable genes used as background to avoid bias.

For each modality, we then identified transcription factors (TFs) associated with each latent factor by testing whether features with non-zero scores were enriched for TF targets. For RNA, we used the CollecTRI regulon database [98] to obtain TF–gene target relationships and, for each TF, compared the scores of its target genes to those of non-target genes using a one-sided Wilcoxon rank-sum test (implemented in scipy.stats.ranksums). For ATAC, we first annotated peaks with TF motifs using gimmemotifs [99] and the JASPAR2026 vertebrates database [100], retaining TF–peak associations with motif scores above 10 as recommended. We then applied the same testing procedure as for RNA, comparing the scores of peaks associated with each TF to those of other peaks using a one-sided Wilcoxon rank-sum test.

## Supporting information

Suppelementary Figures and Tables

## 5 Data availability

All data supporting the findings of this study are already publicly available. **10X PBMC**. We retrieved the 10X Multiome Human PBMC (RNA + ATAC) dataset from https://www.10xgenomics.com/datasets/pbmc-from-a-healthy-donor-no-cell-sorting-10-k-1-standard-2-0-0. **OP Multiome and OP Cite**. We retrieved the Multiome and Cite-seq bone marrow datasets from the Open Problems challenge [34]. The GEO accession number is GSE194122 and the data is available at https://www.ncbi.nlm.nih.gov/geo/query/acc.cgi?acc=GSE194122. **HTA**. We retrieved the scRNA-seq, scATAC-seq and Multiome datasets using the HCATonsil-Data package available at https://github.com/massonix/HCATonsilData/tree/devel. The external human tonsil scRNA-seq dataset [70] is available at ArrayExpress (accession numbers: E-MTAB-8999, E-MTAB-9003, and E-MTAB-9005).

## 6 Code availability

*Package*. The Python package for Champollion is hosted at https://github.com/cantinilab/champollion. It can be installed easily by running pip install champollion-omics. *Reproducibility*. Code to reproduce the experiments and figures is available at https://github.com/cantinilab/champollion_reproducibility.

## 7 Contributions

J.S., G.P., and L.C. designed and planned the study. G.P. and L.C. supervised the study. J.S. and L.C. wrote the paper. G.P. revised the manuscript. J.S. developed the tool and performed all the analyses.

## 8 Acknowledgements

The project leading to this manuscript has received funding from the European Union, European Research Council StG and MULTIview-CELL (101115618, to L.C.). In addition, this work was supported by a government grant managed by the Agence Nationale de la Recherche under the France 2030 program, with the reference numbers ANR-24-EXCI-0001,ANR-24-EXCI-0002, ANR-24-EXCI-0003, ANR-24-EXCI-0004, ANR-24-EXCI-0005 (to L.C.). This work was performed using HPC resources from GENCI–IDRIS (Grant 2021-AD011013214). We acknowledge the help of the HPC Core Facility of the Institut Pasteur and Déborah Philipps for the administrative support. The work of G. Peyré was supported by the French government under the management of Agence Nationale de la Recherche as part of the “Investissements d’avenir” program, reference ANR-19-P3IA-0001 (PRAIRIE 3IA Institute) and by the European Research Council (ERC project WOLF). We also would like to thank Francisco Andrade, Aleksandra Deczkowska, Nicolas Serafini, Rémi Trimbour and Matthieu Najm for helpful discussions on this manuscript.

## Notes

### Competing Interest Statement

The authors have declared no competing interest.

## References

[1] Goodwin, S., McPherson, J. D. & McCombie, W. R. Coming of age: ten years of next-generation sequencing technologies. Nature Reviews Genetics 17, 333–351 (2016). URL 10.1038/nrg.2016.49.

[2] Cusanovich, D. A. et al. Multiplex single-cell profiling of chromatin accessibility by combinatorial cellular indexing. Science 348, 910–914 (2015). URL 10.1126/science.aab1601.

[3] Schwartzman, O. & Tanay, A. Single-cell epigenomics: techniques and emerging applications. Nature Reviews Genetics 16, 716–726 (2015). URL 10.1038/nrg3980.

[4] Bandura, D. R. et al. Mass cytometry: Technique for real time single cell multitarget immunoassay based on inductively coupled plasma time-of-flight mass spectrometry. Analytical Chemistry 81, 6813–6822 (2009). URL 10.1021/ac901049w.

[5] Stoeckius, M. et al. Simultaneous epitope and transcriptome measurement in single cells. Nature Methods 14, 865–868 (2017). URL 10.1038/nmeth.4380.

[6] Swanson, E. et al. Simultaneous trimodal single-cell measurement of transcripts, epitopes, and chromatin accessibility using tea-seq. eLife 10 (2021). URL 10.7554/eLife.63632.

[7] Miao, Z., Humphreys, B. D., McMahon, A. P. & Kim, J. Multi-omics integration in the age of million single-cell data. Nature Reviews Nephrology 17, 710–724 (2021). URL 10.1038/s41581-021-00463-x.

[8] Booeshaghi, A. S., Gao, F. & Pachter, L. Assessing the multimodal tradeoff (2021). URL 10.1101/2021.12.08.471788.

[9] Xu, Y. & McCord, R. P. Diagonal integration of multimodal single-cell data: potential pitfalls and paths forward. Nature Communications 13 (2022). URL 10.1038/s41467-022-31104-x.

[10] Cao, Z.-J. & Gao, G. Multi-omics single-cell data integration and regulatory inference with graph-linked embedding. Nature Biotechnology 40, 1458–1466 (2022). URL 10.1038/s41587-022-01284-4.

[11] Cao, K., Gong, Q., Hong, Y. & Wan, L. A unified computational framework for single-cell data integration with optimal transport. Nature Communications 13 (2022). URL 10.1038/s41467-022-35094-8.

[12] Stuart, T. et al. Comprehensive integration of single-cell data. Cell 177, 1888–1902.e21 (2019). URL 10.1016/j.cell.2019.05.031.

[13] Hao, Y. et al. Dictionary learning for integrative, multimodal and scalable singlecell analysis. Nature Biotechnology 42, 293–304 (2023). URL 10.1038/s41587-023-01767-y.

[14] Gong, B., Zhou, Y. & Purdom, E. Cobolt: integrative analysis of multimodal single-cell sequencing data. Genome Biology 22 (2021). URL 10.1186/s13059-021-02556-z.

[15] Ashuach, T. et al. Multivi: deep generative model for the integration of multimodal data. Nature Methods 20, 1222–1231 (2023). URL 10.1038/s41592-023-01909-9.

[16] He, Z. et al. Mosaic integration and knowledge transfer of single-cell multimodal data with midas. Nature Biotechnology 42, 1594–1605 (2024). URL 10.1038/s41587-023-02040-y.

[17] Du, J.-H., Cai, Z. & Roeder, K. Robust probabilistic modeling for single-cell multimodal mosaic integration and imputation via scvaeit. Proceedings of the National Academy of Sciences 119 (2022). URL 10.1073/pnas.2214414119.

[18] Litinetskaya, A. et al. Integration and querying of multimodal single-cell data with poe-vae (2022). URL 10.1101/2022.03.16.484643.

[19] Zhang, Z. et al. scmomat jointly performs single cell mosaic integration and multi-modal bio-marker detection. Nature Communications 14 (2023). URL 10.1038/s41467-023-36066-2.

[20] Kriebel, A. R. & Welch, J. D. Uinmf performs mosaic integration of single-cell multi-omic datasets using nonnegative matrix factorization. Nature Communications 13 (2022). URL 10.1038/s41467-022-28431-4.

[21] Ghazanfar, S., Guibentif, C. & Marioni, J. C. Stabilized mosaic single-cell data integration using unshared features. Nature Biotechnology 42, 284–292 (2023). URL 10.1038/s41587-023-01766-z.

[22] Dupuy, A. & Galichon, A. Personality traits and the marriage market. Journal of Political Economy 122, 1271–1319 (2014). URL 10.1086/677191.

[23] Galichon, A. & Salanié, B. Cupid’s invisible hand: Social surplus and identification in matching models. The Review of Economic Studies 89, 2600–2629 (2021). URL 10.1093/restud/rdab090.

[24] Carlier, G., Dupuy, A., Galichon, A. & Sun, Y. ¡scp¿sista¡/scp¿: Learning optimal transport costs under sparsity constraints. Communications on Pure and Applied Mathematics 76, 1659–1677 (2022). URL 10.1002/cpa.22047.

[25] Andrade, F., Peyré, G. & Poon, C. Yue, Y. (ed.) Sparsistency for inverse optimal transport. (ed.Yue, Y.) The Twelfth International Conference on Learning Representations (2024). URL https://openreview.net/forum?id=wpXGPCBOTX.

[26] Peyré, G., Cuturi, M. et al. Computational optimal transport: With applications to data science. Foundations and Trends® in Machine Learning 11, 355–607 (2019).

[27] Klein, D. et al. Mapping cells through time and space with moscot. Nature 638, 1065–1075 (2025). URL 10.1038/s41586-024-08453-2.

[28] Demetci, P., Santorella, R., Chakravarthy, M., Sandstede, B. & Singh, R. Scotv2: Single-cell multiomic alignment with disproportionate cell-type representation. Journal of Computational Biology 29, 1213–1228 (2022). URL 10.1089/cmb.2022.0270.

[29] Charlier, B., Feydy, J. & Glaunès, J. Kernel operations on the gpu, with autodiff, without memory overflows. http://www.kernel-operations.io (2018). Accessed: 2018-09-01.

[30] Virshup, I. et al. The scverse project provides a computational ecosystem for single-cell omics data analysis. Nature Biotechnology 41, 604–606 (2023). URL 10.1038/s41587-023-01733-8.

[31] Fu, S. et al. Benchmarking single-cell multi-modal data integrations. Nature Methods 22, 2437–2448 (2025). URL 10.1038/s41592-025-02737-9.

[32] Cao, K., Bai, X., Hong, Y. & Wan, L. Unsupervised topological alignment for single-cell multi-omics integration. Bioinformatics 36, i48–i56 (2020). URL 10.1093/bioinformatics/btaa443.

[33] Singh, R. et al. Aluru, S., Kalyanaraman, A. & Wang, M. (eds) Unsupervised manifold alignment for single-cell multi-omics data. (eds Aluru, S., Kalyanaraman, A. & Wang, M.) Proceedings of the 11th ACM International Conference on Bioinformatics, Computational Biology and Health Informatics, BCB ‘20, 1– 10 (Association for Computing Machinery, New York, NY, USA, 2020). URL 10.1145/3388440.3412410.

[34] Luecken, M. et al. Vanschoren, J. & Yeung, S. (eds) A sandbox for prediction and integration of dna, rna, and proteins in single cells. (eds Vanschoren, J. & Yeung, S.) Proceedings of the Neural Information Processing Systems Track on Datasets and Benchmarks, Vol. 1 (2021). URL https://datasets-benchmarks-proceedings.neurips.cc/paper_files/paper/2021/file/158f3069a435b314a80bdcb024f8e422-Paper-round2.pdf.

[35] van den Berg, P. R., Budnik, B., Slavov, N. & Semrau, S. Dynamic posttranscriptional regulation during embryonic stem cell differentiation (2017). URL 10.1101/123497.

[36] Valk, E., Rudd, C. E. & Schneider, H. Ctla-4 trafficking and surface expression. Trends in Immunology 29, 272–279 (2008). URL 10.1016/j.it.2008.02.011.

[37] Borrego, F., Masilamani, M., Marusina, A. I., Tang, X. & Coligan, J. E. The cd94/nkg2 family of receptors: From molecules and cells to clinical relevance. Immunologic Research 35, 263–278 (2006). URL 10.1385/ir:35:3:263.

[38] Orbelyan, G. A. et al. Human nkg2e is expressed and forms an intracytoplasmic complex with cd94 and dap12. The Journal of Immunology 193, 610–616 (2014). URL 10.4049/jimmunol.1400556.

[39] Walch, M. et al. Cytotoxic cells kill intracellular bacteria through granulysin-mediated delivery of granzymes. Cell 157, 1309–1323 (2014). URL 10.1016/j.cell.2014.03.062.

[40] Srinivasan, S., Zhu, C. & McShan, A. C. Structure, function, and immunomodulation of the cd8 co-receptor. Frontiers in Immunology 15 (2024). URL 10.3389/fimmu.2024.1412513.

[41] Yuan, L. et al. Cd3d is an independent prognostic factor and correlates with immune infiltration in gastric cancer. Frontiers in Oncology 12 (2022). URL 10.3389/fonc.2022.913670.

[42] Schall, T. J., Bacon, K., Toy, K. J. & Goeddel, D. V. Selective attraction of monocytes and t lymphocytes of the memory phenotype by cytokine rantes. Nature 347, 669–671 (1990). URL 10.1038/347669a0.

[43] Topper, M. J. et al. Derivation of cd8+ t cell infiltration potentiators in non-small-cell lung cancer through tumor microenvironment analysis. iScience 26, 107095 (2023). URL 10.1016/j.isci.2023.107095.

[44] Prajapati, K., Perez, C., Rojas, L. B. P., Burke, B. & Guevara-Patino, J.A. Functions of nkg2d in cd8+ t cells: an opportunity for immunotherapy. Cellular and Molecular Immunology 15, 470–479 (2018). URL 10.1038/cmi.2017.161.

[45] Patiño-Lopez, G. et al. Human class-i restricted t cell associated molecule is highly expressed in the cerebellum and is a marker for activated nkt and cd8+ t lymphocytes. Journal of Neuroimmunology 171, 145–155 (2006). URL 10.1016/j.jneuroim.2005.09.017.

[46] Steiner, J. et al. Human cd8+ t cells and nk cells express and secrete s100b upon stimulation. Brain, Behavior, and Immunity 25, 1233–1241 (2011). URL 10.1016/j.bbi.2011.03.015.

[47] Hwang, J. Y. et al. Exploring the expression and function of t cell surface markers identified through cellular indexing of transcriptomes and epitopes by sequencing. Yonsei Medical Journal 65, 544 (2024). URL 10.3349/ymj.2023.0639.

[48] Huse, K. et al. Mechanism of cd79a and cd79b support for igm+ b cell fitness through b cell receptor surface expression. The Journal of Immunology 209, 2042–2053 (2022). URL 10.4049/jimmunol.2200144.

[49] Revilla-i-Domingo, R. et al. The b-cell identity factor pax5 regulates distinct transcriptional programmes in early and late b lymphopoiesis. The EMBO Journal 31, 3130–3146 (2012). URL 10.1038/emboj.2012.155.

[50] Dymecki, S. M., Niederhuber, J. E. & Desiderio, S. V. Specific expression of a tyrosine kinase gene, blk, in b lymphoid cells. Science 247, 332–336 (1990). URL 10.1126/science.2404338.

[51] Pavlasova, G. & Mraz, M. The regulation and function of cd20: an “enigma” of b-cell biology and targeted therapy. Haematologica 105, 1494–1506 (2020). URL 10.3324/haematol.2019.243543.

[52] Mensah, F. F. K. et al. Cd24 expression and b cell maturation shows a novel link with energy metabolism: Potential implications for patients with myalgic encephalomyelitis/chronic fatigue syndrome. Frontiers in Immunology 9 (2018). URL 10.3389/fimmu.2018.02421.

[53] Iaccarino, I. et al. Long non-coding rna linc00926 is a biomarker for naïve b-cells with prognostic value in advanced stage classic hodgkin lymphoma. Haematologica (2025). URL 10.3324/haematol.2025.287524.

[54] Zhang, K. et al. Ramanomics decodes spatial vibrational-molecular architecture and rewiring in aging and repair (2025). URL 10.64898/2025.12.04.692337.

[55] Silverstein, R. L. & Febbraio, M. Cd36, a scavenger receptor involved in immunity, metabolism, angiogenesis, and behavior. Science Signaling 2 (2009). URL 10.1126/scisignal.272re3.

[56] Ferracini, M., Rios, F. J. O., Pecenin, M. & Jancar, S. Clearance of apoptotic cells by macrophages induces regulatory phenotype and involves stimulation of cd36 and platelet-activating factor receptor. Mediators of Inflammation 2013, 1–8 (2013). URL 10.1155/2013/950273.

[57] Chen, X. et al. Identification of fcn1 as a novel macrophage infiltration-associated biomarker for diagnosis of pediatric inflammatory bowel diseases. Journal of Translational Medicine 21 (2023). URL 10.1186/s12967-023-04038-1.

[58] Erbani, J. D. et al. Cd162 is a key e-selectin receptor promoting acute myeloid leukemia chemo-resistance in the bone marrow niche. Blood 134, 907–907 (2019). URL 10.1182/blood-2019-132233.

[59] Erbani, J., Tay, J., Barbier, V.Levesque, J.-P. & Winkler, I. G. Acute myeloid leukemia chemo-resistance is mediated by e-selectin receptor cd162 in bone marrow niches. Frontiers in Cell and Developmental Biology 8 (2020). URL 10.3389/fcell.2020.00668.

[60] Culver-Cochran, A. E. et al. Chemotherapy resistance in acute myeloid leukemia is mediated by a20 suppression of spontaneous necroptosis. Nature Communications 15 (2024). URL 10.1038/s41467-024-53629-z.

[61] Koblish, H. et al. Preclinical characterization of incb053914, a novel pan-pim kinase inhibitor, alone and in combination with anticancer agents, in models of hematologic malignancies. PLOS ONE 13, e0199108 (2018). URL 10.1371/journal.pone.0199108.

[62] Bruynzeel, I., Koopman, G., van der Raaij, L. M., Pals, S. T. & Willemze, R. Cd44 antibody stimulates adhesion of peripheral blood t cells to keratinocytes through the leukocyte function–associated antigen-1/intercellular adhesion molecule-1 pathway. Journal of Investigative Dermatology 100, 424–428 (1993). URL 10.1111/1523-1747.ep12472106.

[63] Guo-Wei, L., Jian-Ping, Q., Xiu-Fang, L. & Yan-Ping, J. Itgb2 fosters the cancerous characteristics of ovarian cancer cells through its role in mitochondrial glycolysis transformation. Aging (2024). URL 10.18632/aging.205529.

[64] Fan, J., Sha, T. & Ma, B. Cancer-derived extracellular vesicle itgb2 promotes the progression of triple-negative breast cancer via the activation of cancer-associated fibroblasts. Global Challenges 9 (2025). URL 10.1002/gch2.202400235.

[65] Rasbach, E. et al. Targeting the tumor cell-intrinsic itgb2 axis inhibits melanoma progression. Molecular Cancer 24 (2025). URL 10.1186/s12943-025-02527-z.

[66] Regev, A. et al. The human cell atlas white paper (2018). URL https://arxiv.org/abs/1810.05192.

[67] Massoni-Badosa, R. et al. An atlas of cells in the human tonsil. Immunity 57, 379–399.e18 (2024). URL 10.1016/j.immuni.2024.01.006.

[68] Moinfar, A. A. & Theis, F. J. Disentangling cellular heterogeneity into interpretable biological factors through structured latent representations (2024). URL 10.1101/2024.11.06.622266.

[69] Luckey, C. J. et al. Memory t and memory b cells share a transcriptional program of self-renewal with long-term hematopoietic stem cells. Proceedings of the National Academy of Sciences 103, 3304–3309 (2006). URL 10.1073/pnas.0511137103.

[70] King, H. W. et al. Single-cell analysis of human b cell maturation predicts how antibody class switching shapes selection dynamics. Science Immunology 6 (2021). URL 10.1126/sciimmunol.abe6291.

[71] Guldenpfennig, C., Teixeiro, E. & Daniels, M. Nf-kb’s contribution to b cell fate decisions. Frontiers in Immunology 14 (2023). URL 10.3389/fimmu.2023.1214095.

[72] Hayden, M. S. & Ghosh, S. Shared principles in nf-κb signaling. Cell 132, 344–362 (2008). URL 10.1016/j.cell.2008.01.020.

[73] Liu, G. et al. Bach2 drives the development and function of group 2 innate lymphoid cells. Science Advances 11 (2025). URL 10.1126/sciadv.ads4323.

[74] Zhang, H. et al. Bach2 attenuates il-2r signaling to control treg homeostasis and tfr development. Cell Reports 35, 109096 (2021). URL 10.1016/j.celrep.2021.109096.

[75] Zwick, D., Vo, M. T., Shim, Y. J., Reijonen, H. & Do, J.-s. Bach2: The future of induced t-regulatory cell therapies. Cells 13, 891 (2024). URL 10.3390/cells13110891.

[76] Montaldo, E. et al. Human rorγt+cd34+ cells are lineage-specified progenitors of group 3 rorγt+ innate lymphoid cells. Immunity 41, 988–1000 (2014). URL 10.1016/j.immuni.2014.11.010.

[77] Hu, X. et al. The transcription factor mef2c restrains microglial overactivation by inhibiting kinase cdk2. Immunity 58, 946–960.e10 (2025). URL 10.1016/j.immuni.2025.02.026.

[78] He, L., Wang, T., Lu, F.-M., Xu, J. & Cong, H.-L. Mef2c protects against the development of atherosclerosis via inhibiting tlr/nf-κb activation. International Journal of Clinical and Experimental Pathology 9, 4179–4187 (2016).

[79] Swiecki, M. & Colonna, M. Unraveling the functions of plasmacytoid dendritic cells during viral infections, autoimmunity, and tolerance. Immunological Reviews 234, 142–162 (2010). URL 10.1111/j.0105-2896.2009.00881.x.

[80] Gilliet, M., Cao, W. & Liu, Y.-J. Plasmacytoid dendritic cells: sensing nucleic acids in viral infection and autoimmune diseases. Nature Reviews Immunology 8, 594–606 (2008). URL 10.1038/nri2358.

[81] Séjourné, T., Peyré, G. & Vialard, F.-X. Unbalanced Optimal Transport, from theory to numerics, 407–471 (Elsevier, 2023). URL 10.1016/bs.hna.2022.11.003.

[82] Mimitou, E. P. et al. Scalable, multimodal profiling of chromatin accessibility, gene expression and protein levels in single cells. Nature Biotechnology 39, 1246–1258 (2021). URL 10.1038/s41587-021-00927-2.

[83] Pass, B. Multi-marginal optimal transport: theory and applications (2014). URL https://arxiv.org/abs/1406.0026.

[84] Cui, H. et al. scgpt: toward building a foundation model for single-cell multiomics using generative ai. Nature Methods 21, 1470–1480 (2024). URL 10.1038/s41592-024-02201-0.

[85] Heimberg, G. et al. A cell atlas foundation model for scalable search of similar human cells. Nature 638, 1085–1094 (2024). URL 10.1038/s41586-024-08411-y.

[86] Bues, J. et al. Single-cell phenomics through integrated imaging and molecular profiling (2025). URL 10.1101/2025.11.28.690954.

[87] Monge, G. Mémoire sur la théorie des déblais et des remblais. Mem. Math. Phys. Acad. Royale Sci. 666–704 (1781).

[88] Kantorovich, L. On the transfer of masses (in Russian). Doklady Akademii Nauk 37, 227–229 (1942).

[89] Cuturi, M. Burges, C., Bottou, L., Welling, M., Ghahramani, Z. & Weinberger, K. (eds) Sinkhorn distances: Lightspeed computation of optimal transport. (eds Burges, C., Bottou, L., Welling, M., Ghahramani, Z. & Weinberger, K.) Advances in Neural Information Processing Systems, Vol. 26 (Curran Associates, Inc., 2013). URL https://proceedings.neurips.cc/paper_files/paper/2013/file/af21d0c97db2e27e13572cbf59eb343d-Paper.pdf.

[90] Hoff, P. D. Lasso, fractional norm and structured sparse estimation using a hadamard product parametrization. Computational Statistics and Data Analysis 115, 186–198 (2017). URL 10.1016/j.csda.2017.06.007.

[91] Paszke, A. et al. Wallach, H. et al. (eds) Pytorch: An imperative style, high-performance deep learning library. (eds Wallach, H. et al.) Advances in Neural Information Processing Systems, Vol. 32 (Curran Associates, Inc., 2019). URL https://proceedings.neurips.cc/paper_files/paper/2019/file/bdbca288fee7f92f2bfa9f7012727740-Paper.pdf.

[92] Kingma, D. P. & Ba, J. Adam: A method for stochastic optimization (2014). URL https://arxiv.org/abs/1412.6980.

[93] Bredikhin, D., Kats, I. & Stegle, O. Muon: multimodal omics analysis framework. Genome Biology 23 (2022). URL 10.1186/s13059-021-02577-8.

[94] Wolf, F. A., Angerer, P. & Theis, F. J. Scanpy: large-scale single-cell gene expression data analysis. Genome Biology 19 (2018). URL 10.1186/s13059-017-1382-0.

[95] Stuart, T., Srivastava, A., Madad, S., Lareau, C. A. & Satija, R. Single-cell chromatin state analysis with signac. Nature Methods 18, 1333–1341 (2021). URL 10.1038/s41592-021-01282-5.

[96] Virtanen, P. et al. Scipy 1.0: fundamental algorithms for scientific computing in python. Nature Methods 17, 261–272 (2020). URL 10.1038/s41592-019-0686-2.

[97] Kolberg, L. et al. g:profiler—interoperable web service for functional enrichment analysis and gene identifier mapping (2023 update). Nucleic Acids Research 51, W207–W212 (2023). URL 10.1093/nar/gkad347.

[98] Müller-Dott, S. et al. Expanding the coverage of regulons from high-confidence prior knowledge for accurate estimation of transcription factor activities. Nucleic Acids Research 51, 10934–10949 (2023). URL 10.1093/nar/gkad841.

[99] Bruse, N. & Heeringen, S. J. v. Gimmemotifs: an analysis framework for transcription factor motif analysis (2018). URL 10.1101/474403.

[100] Ovek Baydar, D. et al. Jaspar 2026: expansion of transcription factor binding profiles and integration of deep learning models. Nucleic Acids Research 54, D184–D193 (2025). URL 10.1093/nar/gkaf1209.

