## Supplementary material for "CHAMPOLLION: Robust Multi-Omics Integration via Inverse Optimal Transport Using Paired Cells": Suppelementary Figures and Tables

| Protein | Source | Term ID | Term name | $p$-value | Term size | Query size | Intersection |
|---|---|---|---|---|---|---|---|
| CD36 | GO:BP | GO:0043277 | apoptotic cell clearance | 3.12e-2 | 8 | 36 | 4 |
| | GO:MF | GO:0005044 | scavenger receptor activity | 4.90e-2 | 3 | 36 | 3 |
| CD162 | GO:BP | GO:0001906 | cell killing | 4.85e-4 | 82 | 90 | 14 |
| | GO:BP | GO:0001909 | leukocyte mediated cytotoxicity | 2.35e-2 | 65 | 90 | 11 |
| CD18 | WP | WP:WP615 | Senescence and autophagy in cancer | 2.27e-2 | 31 | 86 | 8 |

Table 1: **Enriched functional terms identified for selected surface proteins.** Term size denotes the number of genes in the annotated term, query size the number of genes selected as top interactions from the matrix **A**, and intersection the number of overlapping genes between the two groups.

| **CD36** | **CD162** | **CD18** |
|---|---|---|
| VCAN, FCN1, AHSP, MPO, BLVRB, CD36, CA1, LRP1, RHCE, LGALS3, CA3, CES1, ANKRD9, SLC25A39, HBM, CKB, RFLNB, PRTN3, VENTX, AZU1, ICAM1, HEMGN, H1F0, RETN, ID1, TAL1, CCR2, BASP1, CD163, CYP4F3, CEP95, HMBS, FECH, GIHCG, CDKN1C, MOB1B, SPECC1 | GNLY, GZMB, GZMK, CMC1, FOS, HLA-DRA, SPINK2, AC020656.1, AC020916.1, LGALS1, AZU1, DNTT, RPLP0, SERPINF1, ANKRD28, AC103591.3, GZMH, SELPLG, LRRC26, CAPG, CCDC88A, LAMP5, TYMP, TREM1, CES1, CLEC4C, HLA-DRB1, CD52, PAPOLG, CAPN2, FCGR2A, MIR646HG, RFLNB, HLA-F, SYTL1, AC239799.2, PIM1, MIR3945HG, DMPK, PGGT1B, CKB, LINC00623, ZFP36L1, C12orf75, CCL4, ARID5B, KLRC3, CYP20A1, TNFAIP3, VCL, APOC1, ACTB, CACNA1A, GDI1, TCTE3, TNFRSF17, CLNK, FAS, SEMA4B, FAM49A, UGCG, CTSG, S100A12, IFRD1, RNF213-AS1, ZNF544, ANKRD39, PTCRA, FBXL16, ZBTB42, RETN, WDR1, AC253572.2, FBXL5, RAB27A, Z93241.1, EPHB1, AC091271.1, CLEC4E, ARL6IP1, NRP1, NMUR1, MACO1, THAP6, NR1D2, ADRB2, FOXO4, PADI4, GOLGA8B, TOB1, DUSP2, ZNF181, SH2D1A, CEMIP2, LDLRAP1, RAD9A, CSKMT, MKI67, STX18-AS1, PLXNA4 | CD84, AC092279.1, ALOX5AP, PCNA, KBTBD2, AC007262.2, NRIP1, CD74, ITGB2, CA5B, ISCA1, IRF8, PGD, SAMD9L, ZFAND2B, IGSF6, GZMH, AC010168.2, MTIF2, TANGO2, AL512791.2, ZNF548, WDFY2, PIK3R3, MT2A, IGFBP4, ZNF267, CD44, ELANE, TPP2, PEAK1, ZER1, CD86, APBB1, ERN1, SMARCA2, EFHD2, CXCL8, CLN8, CDA, GAPT, ERICH1, LAMP1, KLK1, KLRC1, ZNF583, CCDC71L, ADARB1, CLEC10A, COG3, VAMP1, WAS, TLE4, UBL7-AS1, SNHG7, CD300E, ZNF622, SLA, TRPV2, PGGT1B, IFNG, HIST1H2AD, MPV17L, FBXO33, SPATA2L, RNF19A, LINC01816, PCGF1, ATG14, ZNF493, TUBA1B, ATP10D, CDK13, LINC02352, SPATA13, HMGB2, RBM34, CLIP4, NCR1, DRC3, COQ10B, FCRLB, BEX1, HIST1H3D, UBL3, CCDC84-DT, NIBAN1, ZDHHC1, IRS2, GABARAPL1, DHRSX, SH3GLB1, GPR157, KIAA1109, GNPDA2, SLC7A11, AC099332.1, ATXN7L3B, ATXN1, AL137802.2 |

Table 2: **Top interacting genes for selected surface proteins.** Genes are ordered by decreasing absolute interaction weight in the inferred **A** matrix.

| **scATAC cell type** | **Allowed scRNA cell types** |
|---|---|
| NBC | Naive |
| Activated NBC | Activated |
| MBC | MBC, MBC FCRL4+ |
| GCBC | GC, DZ GC, LZ GC, FCRL2/3high GC, preGC, Cycling |
| PC | Plasmablast, prePB |
| DN | TIM3+ DN |
| CD4 T | CD4+, TfH, TfR, Treg |
| Naive CD4 T | CD4+ NCM |
| CD8 T | CD8+ Cytotoxic |
| Naive CD8 T | CD8+ NCM |
| cycling T | Cycling T |
| NK | NK |
| ILC | ILC |
| Mono/Macro | MAC1, MAC2, MAC3 |
| DC | cDC1 |
| PDC | pDC |
| FDC | FDC |

Table 3: **Allowed correspondences between scATAC-seq and scRNA-seq cell type annotations.** For each scATAC-seq cell type (left column), we list the set of scRNA-seq cell types considered biologically consistent matches. These correspondences were used to assess whether the closest ATAC cell type under the learned cross-modal metric reflects a meaningful biological relationship. Precursor populations were excluded due to ambiguous annotation.

| Latent factor | Source | Term ID | Term name | $p$-value | Term size | Query size | Intersection |
|---|---|---|---|---|---|---|---|
| RNA 42 | GO:BP | GO:0022402 | cell cycle process | 1.15e-11 | 218 | 95 | 31 |
| | GO:BP | GO:0007049 | cell cycle | 4.08e-11 | 299 | 95 | 35 |
| | GO:BP | GO:0000278 | mitotic cell cycle | 6.71e-11 | 168 | 95 | 27 |
| | WP | WP:WP2446 | retinoblastoma gene in cancer | 7.48e-11 | 24 | 95 | 13 |
| | REAC | REAC:R-HSA-1640170 | Cell Cycle | 1.70e-10 | 83 | 95 | 20 |
| | REAC | REAC:R-HSA-69278 | Cell Cycle, Mitotic | 1.06e-9 | 69 | 95 | 18 |
| | GO:BP | GO:1903047 | mitotic cell cycle process | 6.71e-9 | 139 | 95 | 23 |
| | GO:CC | GO:0098687 | chromosomal region | 3.77e-8 | 52 | 95 | 15 |
| | GO:BP | GO:0006259 | DNA metabolic process | 9.87e-8 | 157 | 95 | 23 |
| | GO:BP | GO:0010564 | regulation of cell cycle process | 1.02e-6 | 128 | 95 | 20 |
| RNA 13 | GO:BP | GO:0001775 | cell activation | 5.96e-3 | 477 | 97 | 32 |
| RNA 36 | GO:CC | GO:0009897 | external side of plasma membrane | 3.97e-3 | 200 | 96 | 20 |
| | GO:MF | GO:0038023 | signaling receptor activity | 9.70e-3 | 519 | 96 | 33 |
| | GO:MF | GO:0060089 | molecular transducer activity | 9.70e-3 | 519 | 96 | 33 |
| | KEGG | KEGG:04060 | cytokine-cytokine receptor interaction | 1.53e-2 | 140 | 96 | 16 |
| | GO:MF | GO:0005102 | signaling receptor binding | 2.41e-2 | 459 | 96 | 30 |
| | WP | WP:WP5473 | cytokine cytokine receptor interaction | 2.69e-2 | 128 | 96 | 15 |
| | WP | WP:WP4494 | selective expression of chemokine receptors | 3.57e-2 | 22 | 96 | 7 |

Table 4: **Enriched functional terms associated with selected RNA latent factors.** Term size denotes the number of genes in the annotated term, query size the number of top genes selected in the latent factor gene set, and intersection the number of overlapping genes between the two groups.

| RNA latent factor | ATAC latent factor | TF | RNA $p$-value | ATAC $p$-value |
|---|---|---|---|---|
| RNA 36 | ATAC 34 | REL | 3.62e-3 | 7.19e-19 |
| | | RELA | 7.85e-3 | 8.23e-45 |
| | | NFKB1 | 1.25e-3 | 9.66e-8 |
| | | NFKB2 | 2.88e-4 | 2.45e-22 |
| RNA 42 | ATAC 94 | BACH2 | 1.37e-2 | 3.49e-5 |
| | | FOXS1 | 3.11e-2 | 1.03e-3 |
| | | RORC | 1.95e-2 | 6.14e-4 |
| RNA 35 | ATAC 55 | MEF2C | 2.22e-2 | 3.77e-2 |

Table 5: **Transcription factors shared between selected interacting RNA and ATAC latent factors.** For each latent factor pair, the table reports transcription factors inferred independently from the RNA and ATAC gene sets, restricted to their intersection, together with the corresponding enrichment $p$-values in each modality.

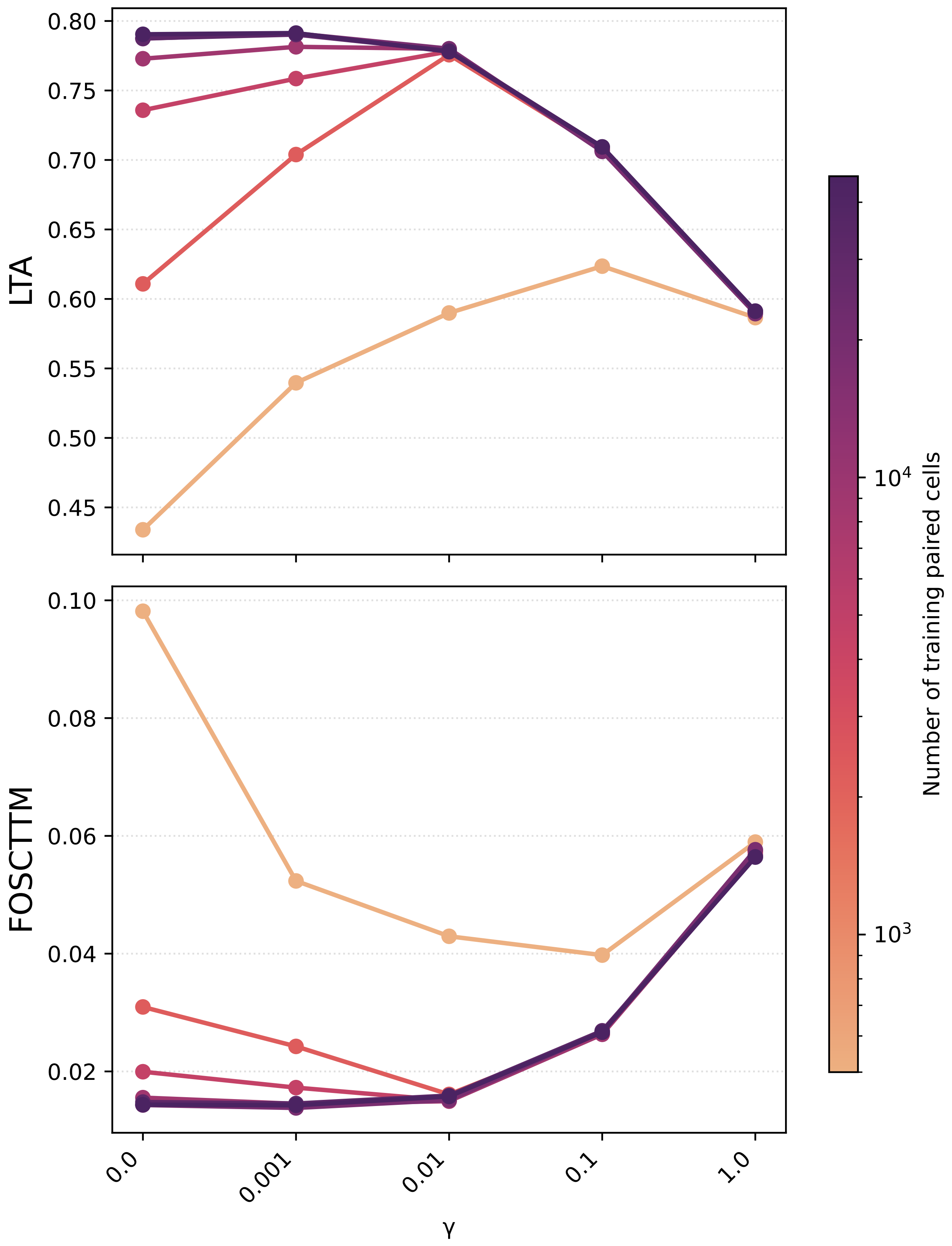


Figure 1: **Impact of gamma on performance.** Performance as a function of the regularization parameter $\gamma$ on the CITE OP benchmark dataset. LTA (top) and FOSCTTM (bottom) are shown for multiple bridge sizes, with each curve corresponding to a different number of paired training cells.

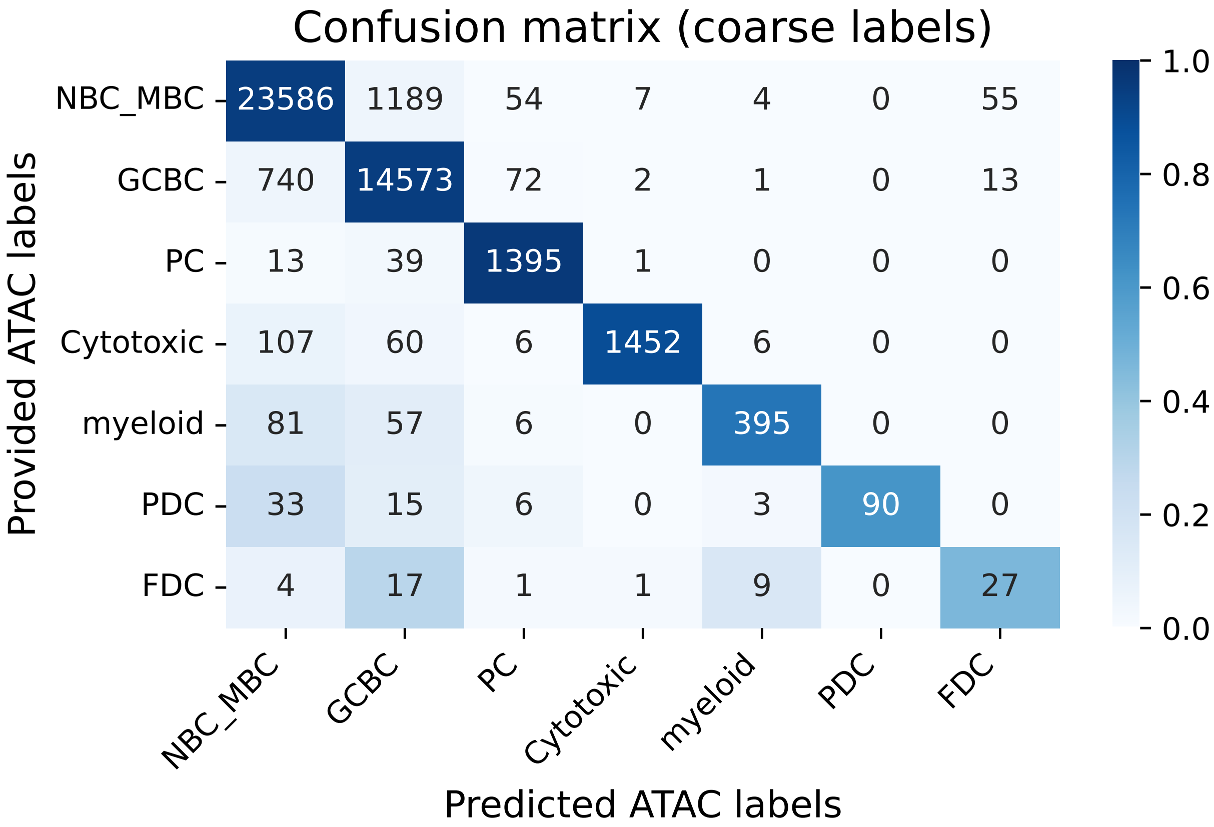


Figure 2: **Coarse label confusion matrix with counts.** Row-wise normalized confusion matrix comparing coarse cell type labels transferred from RNA to ATAC unpaired cells by the dataset authors (rows) with labels inferred by CHAMPOLLION (columns), as in Fig. 4b. Colors indicate row-wise proportions, and numbers denote absolute cell counts.

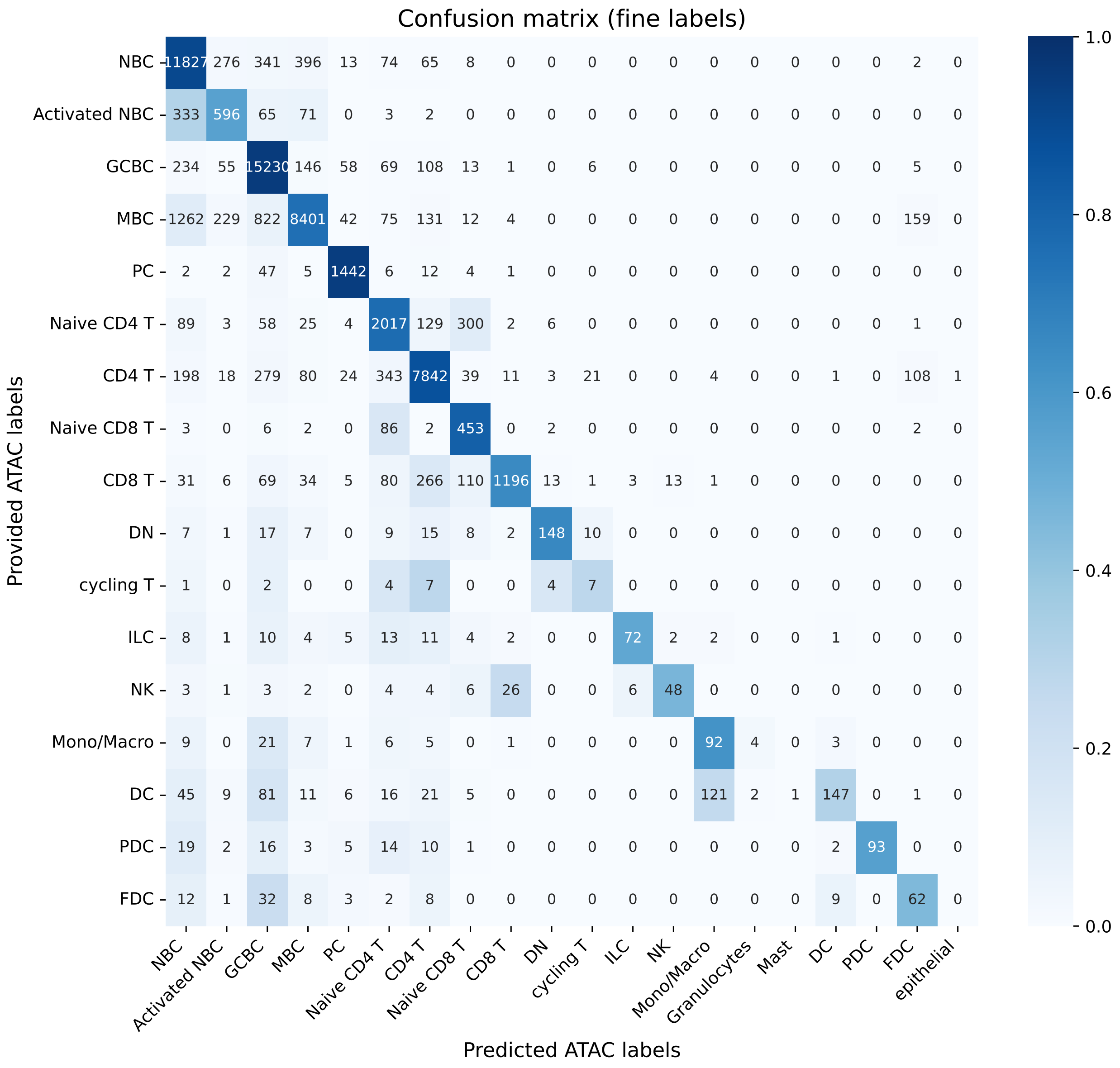


Figure 3: **Fine label confusion matrix with counts.** Row-wise normalized confusion matrix comparing fine cell type labels transferred from RNA to ATAC unpaired cells by the dataset authors (rows) with labels inferred by CHAMPOLLION (columns), as in Fig. 4c. Colors indicate row-wise proportions, and numbers denote absolute cell counts.

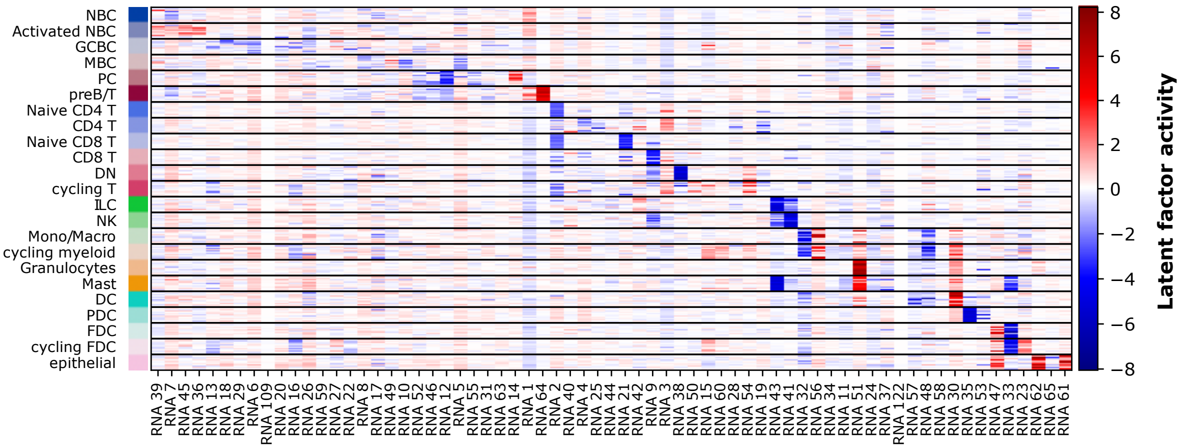


Figure 4: **DRVI latent factors for the RNA modality.** The heatmap illustrates each factor's activity in cells vertically grouped by cell type annotations. Cells are subsampled to have an equal number of cells in each cell type and vanished factors are omitted.

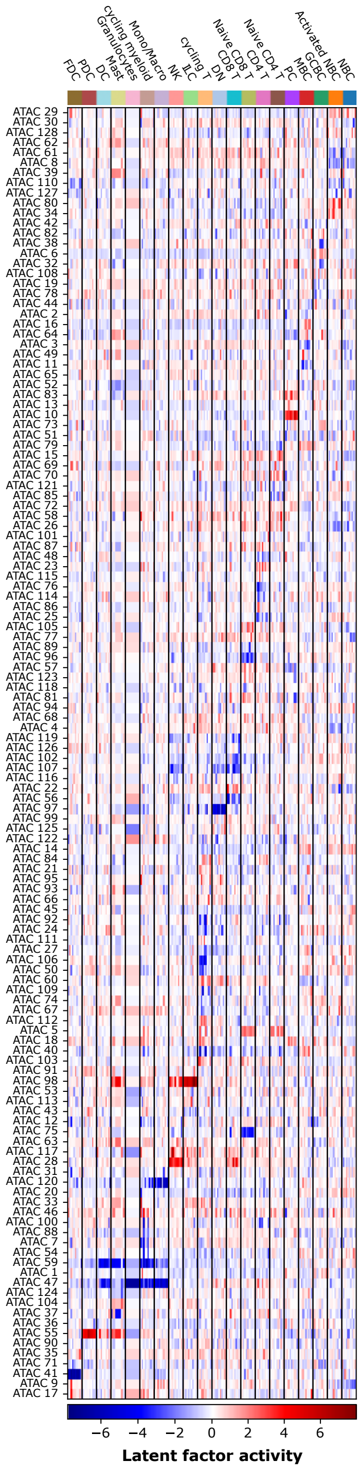


Figure 5: **DRVI latent factors for the ATAC modality.** The heatmap illustrates each factor's activity in cells horizontally grouped by cell type annotations. Cells are subsampled to have an equal number of cells in each cell type and vanished factors are omitted.
